# Protofibril Binding Peptides Recognize and Inhibit Huntingtin Amyloid Formation *in vitro* and *in vivo*

**DOI:** 10.1101/2025.10.21.680574

**Authors:** Kaori Noridomi, J. Mario Isas, Anakha Ajayan, Tristan McPhail, Hui Xu, Anoop Rawat, Christopher Hughes, Natalie Chen, Joshua Lugo, Terry T. Takahashi, Jeannie Chen, Ralf Langen, Richard W. Roberts

## Abstract

In Huntingtons disease, polyglutamine expansion in huntingtin exon 1 (Httex1) results in stepwise misfolding, amyloid formation, and neuronal death. Here we used mRNA display directed evolution to generate peptide ligands targeting Httex1 protofibrils, an early, toxic misfolding intermediate. Two distinct peptide families bind protofibrils, one tryptophan rich and the other glutamine rich, resulting in two predominant peptides HD1 (W-rich) and HD8 (Q-rich). Both peptides bind with high affinity and specificity to the misfolded polyQ structure present in protofibrils, a toxic component that is not recognized by existing huntingtin-directed antibodies. Homo- and heterodimers of HD1 and HD8 bind protofibrils with antibody-like affinity, and potently inhibit aggregation of recombinant and cellular Httex1. The HD8-1 heterodimer can be used like an antibody for immunocytochemistry to identify Httex1 aggregates in transfected cells and in the retina of a Huntingtons disease mouse model system (R6/1). Peptide binding to both *in vitro* and *in vivo* Httex1 validates *in vitro* generated Httex1 protofibrils share the same structural features as Httex1 amyloid from cellular and mammalian disease model systems. Further, HD8-1 protofibril recognition enables direct detection of a pathogenic form of Httex1 (misfolded polyQ) as a disease biomarker. Finally, HD1, HD8, and HD8-1 binding and aggregation inhibition defines the protofibril sites that mediate fibril growth. Overall, our observations support developing protofibril-directed ligands as novel, selective, diagnostics and therapeutics for Huntingtons disease.

**Significance Statement:** Huntingtin protein polyQ misfolding plays a central role in Huntington’s disease pathogenesis. We used mRNA display to discover a new class of huntingtin-directed ligands protofibril-binding peptides. Unlike presently available antibodies, these peptides selectively recognize misfolded polyQ region, enable using toxic huntingtin aggregates as disease biomarkers, and block huntingtin aggregation *in vitro* and *in vivo*. Protofibril binding peptides may thus assist analysis of huntingtin pathology and provide a starting point for designing protein misfolding inhibitors as Huntington’s disease therapeutics.

## Main Text

Huntington’s disease (HD) is a debilitating neurodegenerative disorder caused by an expansion of 36 or more consecutive Gln residues in exon1 (Httex1) of mutant huntingtin (Htt) (1–3). The length of these polyQ expansions is roughly inversely related with disease onset, and biochemical and pathological studies have indicated that increasing Q-lengths promote Htt aggregation into amyloid-like structures (4). In the disease, these inclusions are formed by N-terminal fragments of Htt that are generated by alternative splicing or proteolysis of Htt (5–7). A commonly observed fragment in such aggregates is exon 1 of huntingtin (Httex1), which is sufficient for inducing disease pathology in animal models (8).

As in many amyloid diseases, prevention of aggregation into toxic misfolded structures is considered a potential therapeutic target. Toward this end, it is essential to develop therapeutic molecules that can prevent aggregation and biomarkers that can reliably detect the relevant misfolded species. The aggregation mechanism of Httex1 has been extensively studied (9). Httex1 contains 3 main domains an N-terminal region that has a high helical propensity (N17), the polyQ region containing a Gln repeat of variable length, and a C-terminal proline rich domain (PRD) (Fig. 1A). While the primary driver of Httex1 misfolding is the beta-sheet formation of the expanded polyQ region, the N- and C-terminal domains have strong modulatory roles. The N17 promotes aggregation and is responsible for the formation of helical structures that can be detected early during aggregation *in vitro* (10–12). Subsequently, small beta-sheet containing oligomers and protofibrils can be detected, which can act as seeds that recruit native Httex1 into fibrillar structures (11). The beta sheet present in these structures is in the polyQ region. Cell studies suggest that early oligomers and protofibrils are more toxic than the more bundled fibrils (13–15). Moreover, protofibrils are much more potent seeds than more mature, bundled fibrils (13). Thus, binders that recognize these early misfolding intermediates might not only find utility as biomarkers for toxic misfolding intermediates, but they might also be able to interfere with the aggregation pathway.

**Figure. 1.**
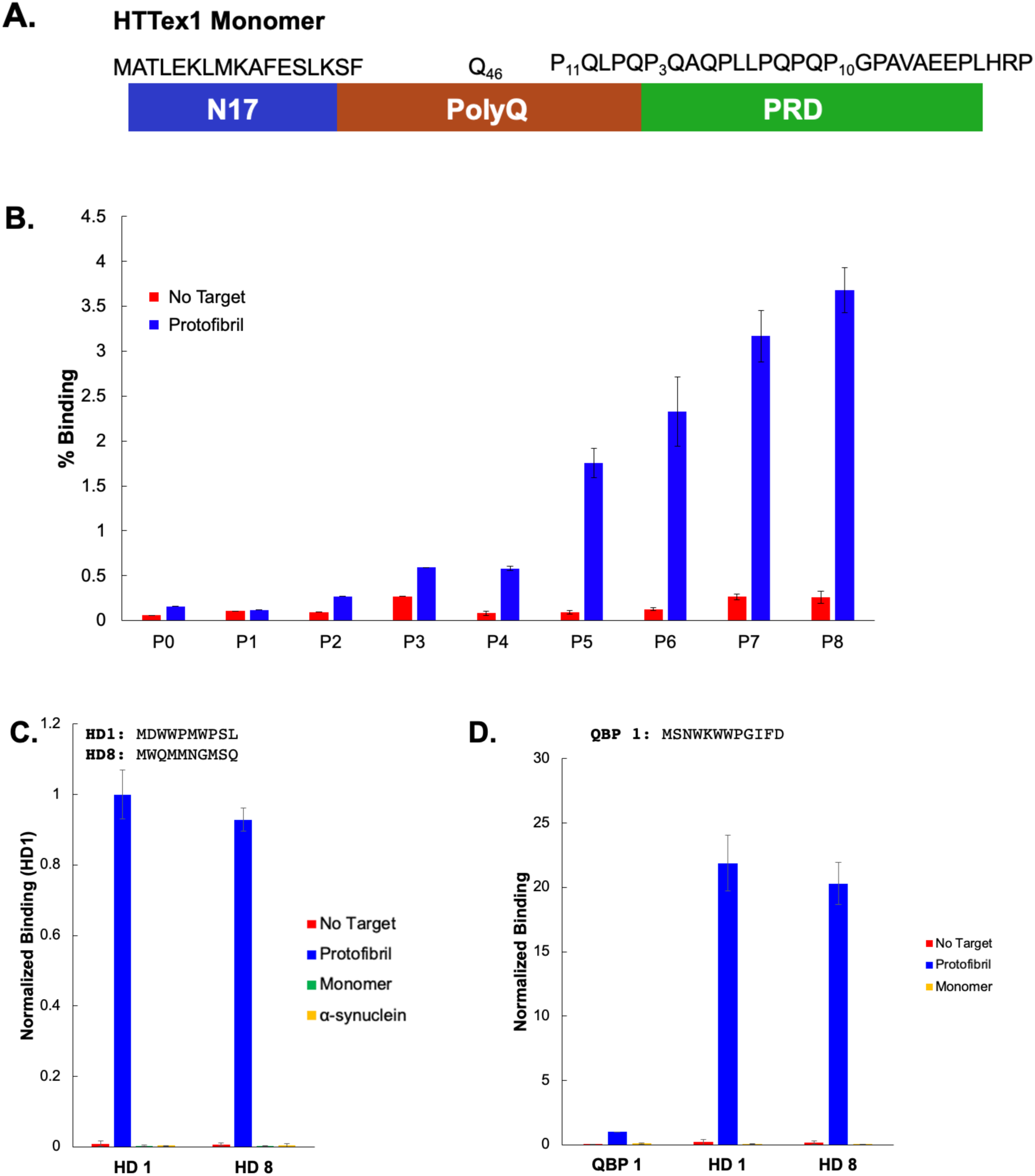
mRNA Display selection to identify HTTex1 protofibril-binding peptides. *(A)* Monomeric Httex1 contains three sequence motifs: i) N17, ii) the polyglutamine repeat (polyQ), and iii) the proline rich domain (PRD). *(B)* Binding and selection of Httex1Q46 protofibril-binding peptides using a naïve MX_9_ starting library, from pool 0 through pool 8. *(C)* HD1 and HD8 peptide sequence and binding to immobilized Httex1(Q46) protofibrils (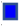), HTTex1(Q46) monomer (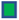), α-synuclein amyloid (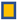), and streptavidin agarose beads without target (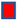) measured by radioactive pulldown. *(D)* Relative affinity of the HD1, HD8, and the poly glutamine binding peptide QBP1 (MSNWKWWPGIFD) (17). Error bars represent the standard deviation of three or more experiments.

The ability to make structurally defined intermediates opens the possibility to directly target protofibrils in a screen for new ligands. Here, we used mRNA display to find novel peptide ligands that bind protofibrils. This technique was chosen for three reasons. First, the high sequence diversity present in mRNA display methodologies has been shown to yield high affinity ligands, typically in the nanomolar range (16). Second, unlike antibody generation, there is no need for the target to be stable for extended period *in vivo*, as the targets can be bound to beads and screened for binders immediately and third, attempts to raise antibodies against misfolded beta-sheet polyQ structures have so far been unsuccessful, generally resulting in ligands that recognize linear or simple epitopes such as the N17, linear polyQ, or the PRD. This work resulted in two chemically distinct peptides that appear to bind different, partially overlapping sites on the protofibrils. These peptides provide a starting framework for developing new diagnostics (e.g., via bioassay, PET labeling) and for generating molecular tools that inhibit fibril formation.

## Results

### mRNA Display Targeting a Httex1 protofibrils

A major focus of our work is developing high affinity peptide ligands that selectively target Httex1 protofibrils. To do this, we prepared sparsely labeled biotinylated Httex1(Q46) protofibrils containing 95% unmodified Httex1(Q46) and 5% biotinylated Httex1(Q46) and immobilized them on streptavidin resin (see Materials and Methods and Table S1). Protofibrils prepared in this manner are relatively stable and suitable as a selection target (13). We targeted the protofibrils via mRNA display using a nine residue random library (X_9_) to perform successive rounds of selection and directed evolution. The naïve starting library contained ∼1 x 10^12^ individual sequences, around two-fold higher than the expected theoretical complexity of the X_9_ pool (20^9^ = 5 x 10^11^ sequences), indicating that most of the possible nine-residue sequences were represented. Binding over background is seen in pool 2 and increased until pool 8 (Fig. 1C). The enriched cDNA libraries from pools six through eight were subjected to high throughput sequencing. These peptides fell into two main groups, one containing one or more tryptophan residues and no glutamines (W-rich) and reads containing one or more glutamines (Q-rich). Testing individual sequences, the two highest abundance peptides, HD1 (W-rich) and HD8 (Q-rich) were identified.

### HD1 and HD8 Binding Specificity

We tested the binding specificity of HD1 and HD8 using radioactive pull-down experiments with immobilized protofibrils. Both HD1 and HD8 show strong pull-down using immobilized protofibrils and essentially no binding to beads loaded with monomeric Httex1Q46, or α-synuclein fibrils (Fig. 1 C), indicating that they are specific for the structural form present in huntingtin protofibrils. We next compared the affinity and specificity of HD1 and HD8 with the previously reported QBP1 peptide (17) that was identified in phage display screens targeting aggregated polyQ (Fig. 1D). While QBP1 binds Httex1 protofibrils with good selectivity, both HD1 and HD8 show ∼20-fold higher pulldown efficiency. Taken together, these observations differentiate HD1 and HD8 binding from both the QBP1 polyQ binding peptide as well as fibril binding dyes such as thioflavin T, which binds to many types of amyloid fibrils.

### Pool analysis defines peptide-protofibril recognition

The abundant round 8 sequences (>500 reads) (Supp. Table 2) can be divided into two groups with approximately 200 sequences each: 1) W-rich sequences containing no glutamines and 2) Q-rich sequences (Fig. 2). Within each group, patterns of W or Q residues appear at regular intervals and the most abundant examples were binned to identify conserved residues in each sequence context. The most abundant W-rich patterns are 1) HD1-like peptides (W3, W4, W7), 2) peptides containing W4 and W7, and 3) peptides containing W4, W7 and W10 (Fig. 2A). These W-rich sequences show similar conservation patterns: 1) D at position 2, 2) W or aromatic at position 3, 3) P at positions 5 and 8, 4) M at position 6, and L at position 10. The pattern of W residues at positions 3, 4, 7, and 10 is consistent with HD1 peptides forming either an α-helix or a polyproline type II helix. However, proline at positions 5 and 8 highly disfavors α-helix formation (18) but is compatible with a polyproline type II helix.

**Figure. 2.**
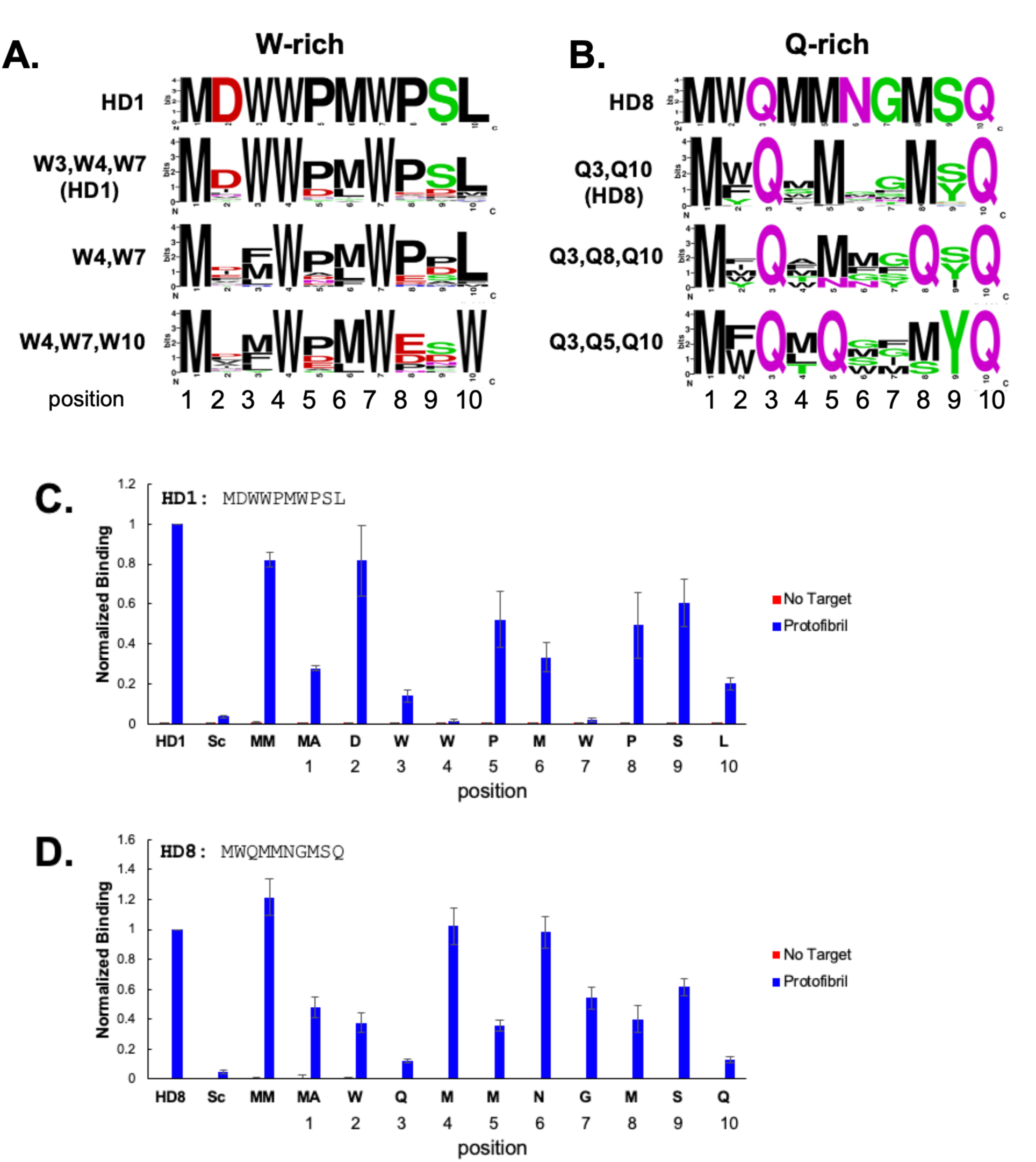
Sequence alignment, conservation, and alanine scanning mutagenesis for protofibril binding peptides. *(A)* Sequence logo representation of W-rich pool 8 sequences binned by tryptophan content i) W3,W4, W7 (HD1-like), ii) W4, W7, and iii) W4, W7, and W10 (56, 57). *(B)* Sequence logo representation of Q-rich pool 8 sequences binned by Q content: i) Q3, Q10 (HD8-like) ii) Q3, Q8, Q10, and iii) and Q3, Q5, Q10 (56, 57). Alanine scanning mutagenesis of *(C)* HD1 and *(D)* HD8 assayed by radioactive pulldown. The normalized binding for single mutations to alanine at each position is shown. The N terminal methionine was probed by adding an additional methionine (MM) and mutating the first position to A (MA). Binding of the scrambled versions of HD1 and HD8 (Sc = Scrambled) is shown.

For the Q-rich sequences, significantly more sequence diversity is observed in the round 8 pool (Fig. 2B, S1). First, although huntingtin aggregation is driven by the polyQ domain, none of the pool 8 sequences show polyQ tracts, with ∼1% of sequences containing either QQ dipeptide or QQQ tripeptide steps (Supp. Table 2). Patterning is seen by grouping the Q-rich peptides by Q position. HD8 and HD8-like peptides (Fig. 2B) show conservation of Q or M at positions 3, 5, 8, and 10. Many sequences contain the MWQ or MFQ tripeptide at the N-terminus and M-S/Y-Q at residues 9 and 10, along with a region of low conservation at position 6 and 7. In these alignments, it appears that Q and M can be swapped while retaining function. The spacing of the Q/M residues at i, i + 2, and i + 4 is compatible with a β-sheet formation and could enable HD8 to bind polyQ aggregates similarly to the interdigitated Q interactions in Orb2 (19). Interestingly, when sequences are binned by the position of the Q residues, there is almost perfect conservation of methionine at the other positions. For example, in the 35 HD8-like sequences (Q3, Q10) there is almost perfect conservation of methionine at positions five and eight (M5, M8). The M and Q interchangeability indicates methionine can substitute for glutamine in the Q-rich binders.

To better understand peptide protofibril recognition, we tested Q-repeat containing peptides (Q7, Q10, Q15, and Q20) for protofibril binding. While we did not observe polyQ containing peptides in the selected pools, peptides containing Q-tracts do bind to Httex1 protofibrils with high affinity (Fig S2). Q7 and Q10 peptides bind over background but bind at lower levels than HD1. Peptides containing Q15 or Q20 bind better than HD1 and there is a trend of increasing protofibril binding as the Q-tract is elongated.

Within these two peptide families, some amino acids increase or decrease in abundance in the final pool 8 vs. the naïve pool 0 libraries (Fig. S3 A, B), indicating residues under positive or negative selection pressure (20). For W-rich peptides, we observe that tryptophan, phenylalanine, methionine, proline, and aspartic acid are enriched, while polar residues are disfavored. For Q-rich sequences, Q, M, and aromatic residues are enriched, while charged and polar residues are under negative selection pressure. Somewhat surprisingly, positively charged residues are almost entirely excluded from both the W-rich and Q-rich sequences. R, H, and K residues are reduced by more than 10-fold in the final pool relative to their initial abundance indicating that positively charged amino acids are highly disfavored for protofibril binding.

We next examined the structure-activity relationship of HD1 and HD8 using alanine scanning mutagenesis (21). For HD1, W3A, W4A, and W7A point mutations decrease the pulldown efficiency by more than 90% (Fig. 2 C.). Ranking the impact for remaining residues (biggest to smallest) gives L10A, M1A, M6A, P8A, P5A, S9A, while D2A has little effect on binding. For the peptides containing W3, W4, and W7, this is the same rank order of conservation seen in the sequence alignment (Fig. 2A). For HD8, Q3A and Q10A both show ∼90% binding reduction and alanine mutations W2A, M5A, M8A, M1A, G7A, S9A, show binding reductions, while M4A and N6A are neutral. The sequence conservation (Fig. 2B) shows a similar ordering (M5 = M8 > W2 = S9 > G7) and positions 4 and 6 show little conservation. Scrambled versions of HD1 (Sc) and HD8 show very little binding. These data identify residues that play an important role in protofibril recognition, indicate that peptide-protofibril binding is specific and sequence dependent, and that aromatic, hydrophobic, and polar residues play important roles in the peptide-fibril interface.

### Protofibril binding and stoichiometry

We next explored the protofibril-peptide interactions using EPR spectroscopy (Fig. 3). Spin-labeled HD1 (Fig. 3A) and HD8 (Fig. 3B) peptides were added to protofibrils. In both cases, the decreased peptide mobility upon protofibril binding resulted in a substantially reduced EPR amplitude, ∼three-fold for HD1 (Fig. 3A) and ∼10-fold for HD8 (Fig. 3B). Amplifying the bound-state signal (blue dashed line) indicates that both peptides undergo distinct and different changes in line shape, with HD8 showing larger changes than HD1. Following our established protocol (22), we used these amplitude changes to determine the fraction of bound and free peptide at various concentrations of added HD1 or HD8 (8, 10 and 12 µM), thereby determining the Httex1/peptide stoichiometry (Fig. 3 C). Those data indicate that there is 1:2.8 ratio for HD1 to Httex1 binding and that HD8 binding is nearly stoichiometric, giving 1:1.2 HD8 binding per Httex1 monomer. The HD1/Httex1 stoichiometry is maximal for fresh protofibrils, and it decreases as the protofibrils mature (see below). Taken together, the sequence, spectroscopic, and kinetic differences observed between HD1 and HD8 argue these peptides identify two different protofibril recognition motifs.

**Figure. 3.**
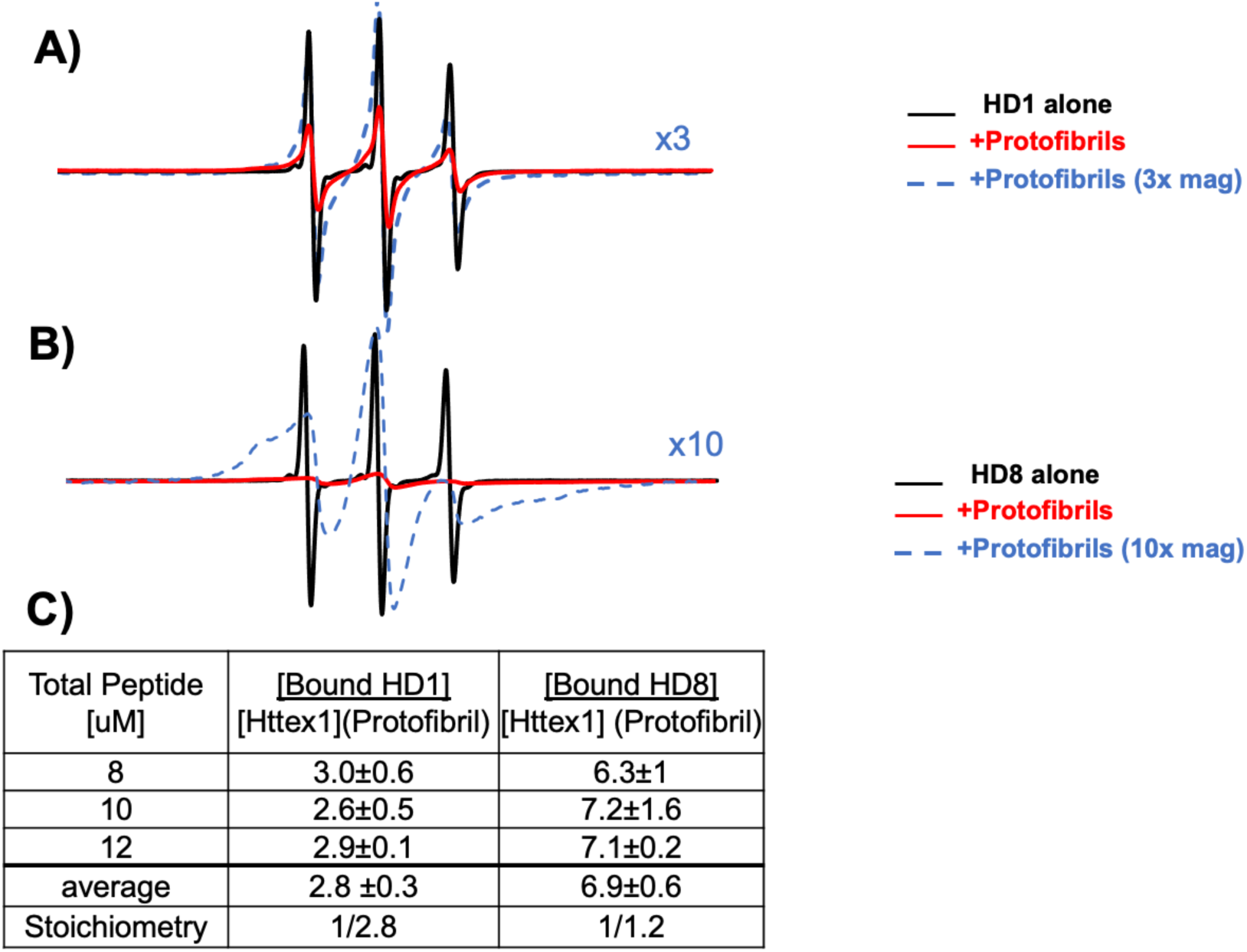
HD1 and HD8 protofibril binding and stoichiometry measured by measured by EPR spectroscopy. The EPR spectra for nitroxide spin-labeled *(A)* HD1 and *(B)* HD8 peptides was measured in free solution (black) and in the presence of excess protofibrils (red). The bound spectra are also shown amplified 3X (HD1) or 10X (HD8) to estimate the amplitude loss and demonstrate changes in EPR line-shape (dashed blue). Spin-labeled HD1 and HD8 peptides show strong binding-dependent decreases in EPR amplitude in the presence of protofibrils. *(C)* Binding and stoichiometry of HD1 and HD8 with Httex1Q46 protofibrils measured by EPR spectroscopy. The moles of bound peptide were measured peptide concentrations of 8, 10, and 12 μM in the presence of 8 μM protofibril target essentially as previously described (22). HD1 shows a stoichiometry of 1 mole of peptide per 2.8 moles of protofibril Httex1, while HD8 shows binding of 1 mole of peptide to 1.2 moles of protofibril Httex1.

### HD1 and HD8 both recognize the polyQ structure in protofibrils

We next used EPR spectroscopy to identify the regions within Httex1 recognized by HD1 and HD8 (Fig. S4). To do this, we prepared Httex1 protofibrils that lack either the PRD (Httex1ΔPRD) or that lack the N17 domain (Httex1ΔN17). Both HD1 and HD8 showed amplitude reduction upon binding to Httex1ΔPRD protofibrils, indicating that the PRD is not essential for binding (Fig. S4 A). Further, HD1 and HD8 also bind protofibrils made from a protein with deleted N17 (Httex1ΔN17; Fig. S4 B, *Cf.* Fig 1A). Since neither the PRD nor the N17 sequences are essential for binding, we conclude that both HD1 and HD8 bind and recognize misfolded polyQ structure present within the protofibrils.

### HD1 and HD8 show little binding to bundled fibrils

The HD peptides were selected using early protofibril aggregates, raising the question of whether they bind bundled fibrils. To test the bundled species, we repeated the EPR experiments of spin labeled HD1 and HD8 peptides in the presence or absence of bundled fibrils (Fig. S5). The addition of bundled fibrils had negligible effects on the EPR spectra of HD1, indicating that there is little interaction. The HD8 EPR signal shows ∼15% amplitude reduction indicating that HD8 peptide/Httex1 binding stoichiometry is reduced around 6-fold in the fibrils compared to the protofibril.

### Homo- and Hetero-Dimers of HD1 and HD8 peptides bind protofibrils with greatly improved affinity

Based on the binding stoichiometry, protofibril formed from multiple Httex1 should contain several binding sites for HD1 and HD8. Further, the HD1 and HD8 binding stoichiometry indicated that they recognize different sites on the protofibril. To test this, we constructed different HD1 and HD8 homodimers and heterodimers containing flexible hydrophilic spacers of zero (SP0) to ten (SP10) amino acids (Fig. 4 A., B. Sup. Fig. 3). We reasoned that such dimers might have improved affinity vs. the parent monomers because both HD1 and HD8 bind the same domain (polyQ region) and because each peptide has relatively high stoichiometry relative to Httex1.

**Fig. 4.**
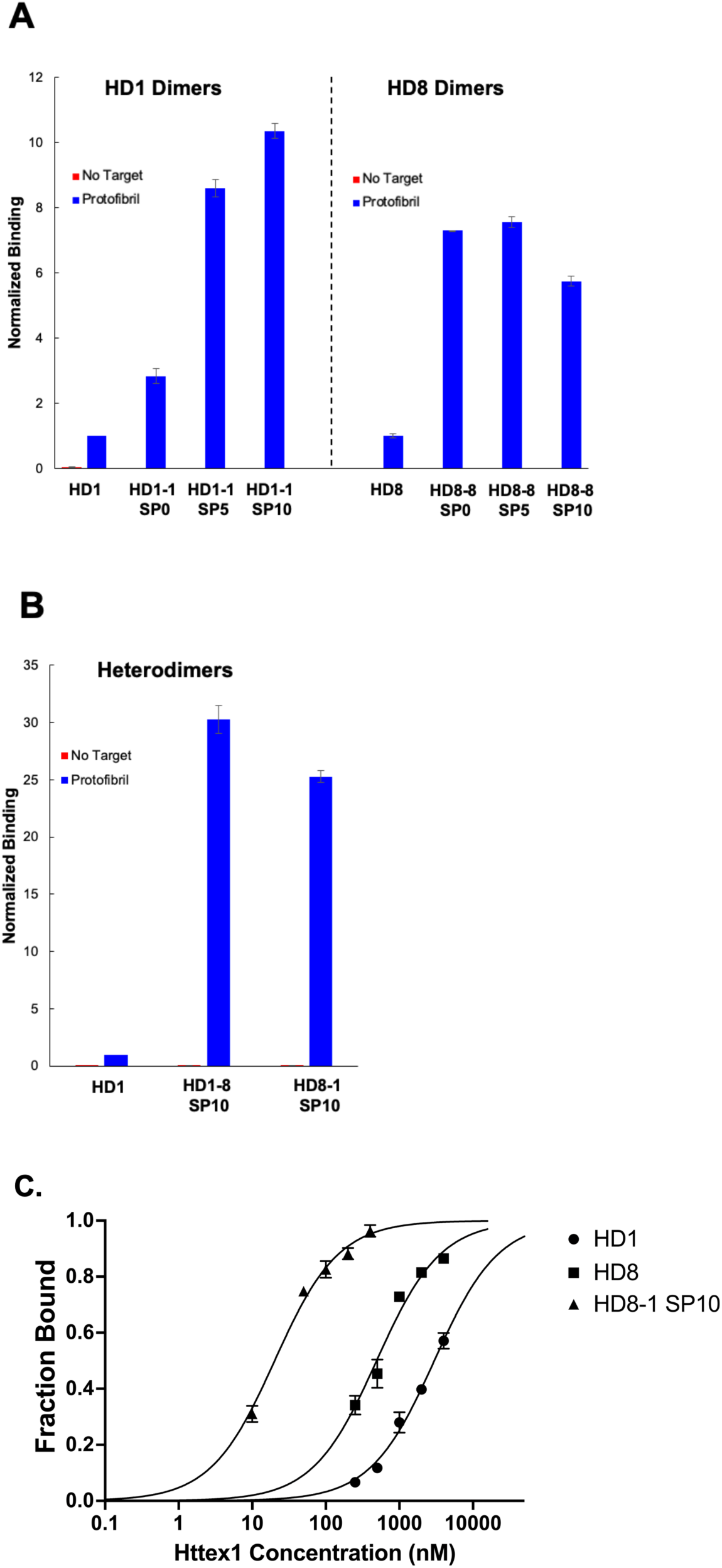
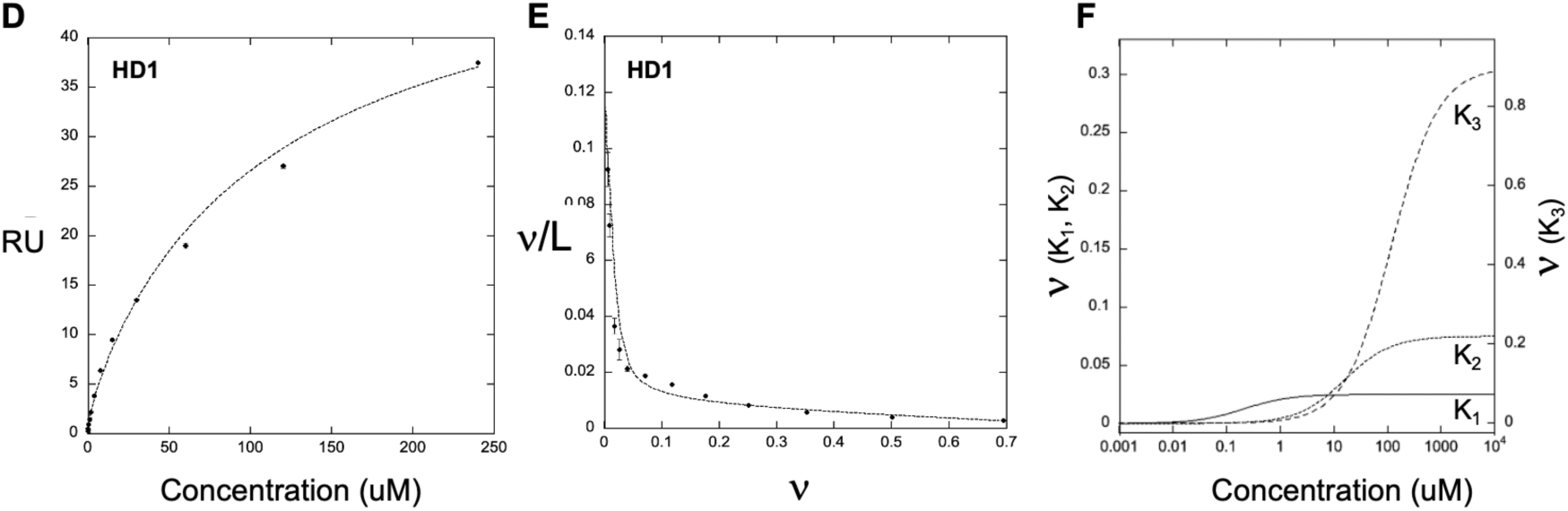
Binding affinity and binding site analysis of HD1, HD8, and homo- and heterodimers of HD1 and HD8 to Httex1Q46 protofibrils. *(A)* Normalized binding of HD1 and HD8 tandem homodimers with no spacer (SP0) a five-residue spacer (SP5) or a ten-residue spacer (SP10). *(B)* Normalized binding of HD1, HD8-1 SP10 heterodimer, and HD1-8 SP10 heterodimer peptides. *(C)* Binding for HD1, HD8, and HD8-1 SP10 peptides measured by radioactive pulldown with Httex1 protofibrils. *(D)* Steady-state binding of HD1 to immobilized protofibrils analyzed by SPR. *(E)* Scatchard analysis of HD1 binding to immobilized Httex1 protofibrils. *(F)* Binding isotherms and amplitudes for high (K_1_) medium (K_2_) and low (K_3_) affinity binding sites for HD1 with Httex1 protofibrils. For plots *E* and *F*, ν = 1 corresponds to a stoichiometry of one site per three Httex1 monomers in the protofibril.

Dimers, trimers, and tetramers show improved binding compared to the individual monomers and affinity also depends on the length of the flexible spacer. For the HD1-1 dimer, a spacer of 10 residues gives ∼10-fold increase in binding compared to HD1. As the spacer is decreased, the affinity decreases and a spacerless dimer showed only 2-fold improvement compared to HD1 alone (Fig. 4 A., Fig. S6). For HD8, a spacerless dimer shows ∼7-fold binding improvement and maximal binding is observed for a four or five amino acid spacer. Heterodimers containing HD1 and HD8 (Fig. 4 B) both bind with markedly higher affinity than either homodimer and give optimal affinity with a long 10 residue spacer. Indeed, HD1-8 SP10 and HD-8-1 SP10 show 25- to 30-fold improved binding compared with HD1.

Along with the polyQ specificity and stoichiometry data, these multimer data indicate that each protofibril has multiple HD1 and HD8 sites and that 10 amino acid linkers enable bi-, tri, and tetradentate interactions within one particle. Further, the fact HD8 and HD1 dimers give strong interactions indicates there are HD8 sites proximal to both the N- and C-terminus of HD1 and HD1 sites proximal to the N and C-terminus of HD8. For HD8, dimers must be able to interact in an almost continuous way, since even dimers with no spacer have improved binding. For HD1 homo- and heterodimers, a 5 residue spacer is sufficient for bidentate binding, indicating the distance between HD1 sites and between HD1/HD8 sites is less than 17Å, the end to end distance for an extended 5 residue chain.

### Affinity of HD1, HD8, and HD8-1 SP10

We next measured the binding affinity of HD1, HD8, and HD8-1 SP10 dimer using radioactive pull-down experiments (Fig. 5) (see Materials and Methods). To do this, we measured radioactive peptide binding to increasing amounts of immobilized protofibrils, generating binding isotherms (Fig. 5 A., B.). All three peptides are slow to come to equilibrium, with HD8-containing peptides taking ∼24 hours to reach steady state. If there were one binding site per Httex1 monomer in the protofibrils, this approach would give HD1 K_d_ = 2.9 ± 0.1 uM, HD8 K_d_ = 490 nM and HD8-1 SP10 dimer as K_d_ = 20 nM. To get the binding constant for each site, we need to divide these values by the stoichiometry (Fig. 3C). For HD1 and HD8-1 SP10, this gives K_d_ = 1.0 uM and 7.1 nM, respectively and for HD8 K_d_ = 410 nM. We also knew that the HD1/Httex1 ratio decreases over time, so we sought to perform binding analysis via SPR where we could get information on both the stoichiometry and affinity in the same experiment.

**Fig. 5.**
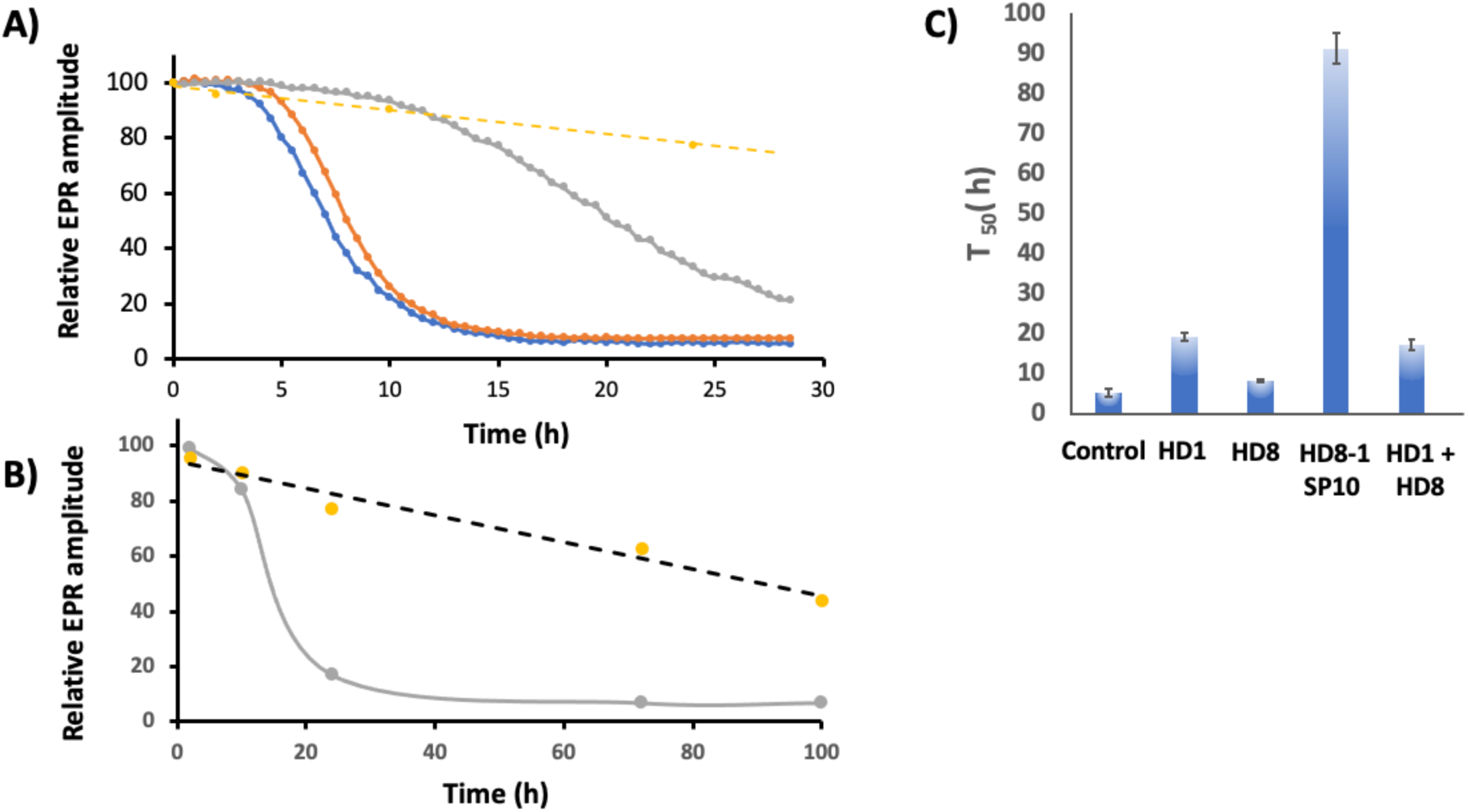
Inhibition of Httex1(Q_46_) aggregation monitored by EPR spectroscopy. Nitroxide labeled Httex1 (Httex1(Q_46_)-35R1) shows a time dependent decrease in amplitude as the protein aggregates. *(A)* representative Httex1 aggregation kinetics in the presence of buffer alone (blue), HD1 (gray), HD8 (orange), and HD8-1 SP10 (yellow) over a 30 hour period. *(B)* Httex1 aggregation kinetics for the HD8-1 SP10 (dashed line) and an HD8 + HD1 admixture (gray) measured over a longer 100 hour time period. *(C)* Summary of the aggregation half-lives (t_50)_) from triplicate kinetics showing the relative inhibition properties of HD1, HD8, HD1 + HD8, and the HD8-1 SP10 peptides.

### Scatchard Analysis Indicates Protofibrils Have More than One Type of HD1 Site

Scatchard analysis enables determining both the number and affinity of ligand binding sites on a particular target (23). We performed Scatchard analysis using surface plasmon resonance steady state HD1-protofibril binding under conditions where the peptide was in vast molar excess to the protofibril (Fig. 4 D,E). HD1 binding is fully reversible to immobilized protofibrils under these conditions while HD8 containing peptides (HD8, HD8-1 SP10) cannot be analyzed this way because they accumulate on the chip over multiple runs. HD1 gives an apparent midpoint at K_d_ ∼90 uM peptide, an unexpected result based on the radioactive binding isotherms (Fig. 4C) and because peptides with weak binding (K_d_ > 1 uM) generally do not show efficient pulldown in radioactive mRNA display format due to the high dissociation rate constants (24). However, the steady state binding is not well fit by a single binding isotherm at both low and high peptide concentrations. Scatchard analysis for HD1 binding provides a route to resolve this issue. When there is a single type of binding site, the Scatchard plot is straight and the slope is –(1/K_eq_), the microscopic K_d_ value. HD1 binding protofibrils produces a highly curved plot indicating there are two or more different kinds of microscopic HD1 binding sites (Fig. 4 E) (25, 26). The SPR and Scatchard analysis supports a three-site model, giving high affinity (K_1_ = 210 nM; 2.5%), intermediate affinity (K_2_ = 16 uM; 7.5%), and low affinity (K_3_ = 120 uM; 90%) HD1 binding sites on the protofibril (Fig. 4. E, F) (data fit as in (25) and (26) see Materials and Methods). In this model, only ∼10% of HD1 binding sites are high or medium affinity (Fig. 4. F) a stoichiometry of one such site per 30 Httex1 monomers. The presence of high affinity HD1 sites also supports HD1 being the most abundant peptide in the round 8 pool and showing higher affinity than HD8 (Fig. 1D,E).

### Competition between HD1, HD8, and dimer peptides

We next probed the peptide-protofibril binding using competition experiments (Supp. Fig. 5), measuring radioactive binding for HD1, HD8, HD1-1 SP10 and HD1-8 SP10 in the presence of increasing amounts of synthetic competitor peptide. Each of the four peptides shows efficient self-competition with synthetic peptide and at high concentration, the synthetic peptide can block more than 95% of the identical radioactive peptide binding. This competition is consistent with the synthetic peptides being identical in sequence and function to the peptide fusions generated in the selection by *in vitro* translation. The competition data provide further support for there being a small proportion of high affinity HD1 sites. Synthetic HD1 (10uM) only reduces HD8 binding by ∼40%, but synthetic HD8 (10 uM) blocks ∼95% HD1 binding. These observations support the Scatchard analysis showing that most HD1 sites are low affinity, arguing that only the high and intermediate affinity sites can functionally compete with HD8 binding.

### HD1 and HD8-1 SP10 Inhibit Httex1 Aggregation

We next evaluated HD1, HD8, and HD8/HD1 dimer peptides ability to alter Httex1 aggregation, using a previously established EPR-based aggregation readout (11). This readout monitors the mobility changes that occur in the polyQ region (residue 35) as the dynamic structure of the monomer is converted into a more rigid β-sheet structure in the fibril. Aggregation in the absence of HD peptides is rapid with a t_50_ of ∼ 5 hours (Fig. 5A, blue trace). Equimolar concentrations of HD8/Httex1 slow aggregation somewhat, giving a t_50_ ∼ 6 hours. On the other hand, equimolar HD1 dramatically slows aggregation by 3.8-fold (t_50_ of ∼19 hours). Further, equimolar HD8-1 SP10 dimer shows an even more pronounced 18-fold reduction with a t_50_ of ∼90 hours (Fig. 5B,C). This reduction is not a consequence of simply having both peptides present, as the admixture of HD1 and HD8 gives similar kinetics to HD1 alone (Fig. 5C). Thus, peptides containing the HD1 sequence inhibit Httex1 aggregation and increasing the affinity (by adding the HD8 sequence) further inhibits the aggregation process. The strong inhibition observed for both HD1 and HD8-1 SP10 indicates that Httex1 misfolding is slowed dramatically by occupying relatively few HD1 high and intermediate affinity binding sites (one site per 30 Httex1 monomers).

### HD8-HD1 recognizes Httex1 Aggregates and Colocalizes with PHP1 in Fixed Cells

A key question is whether the structural elements on in vitro synthesized protofibrils is present in misfolded Httex1 formed in cells. To address this question, HEK293 cells were transfected with a Httex1(Q72) expression construct that readily forms large aggregates after 24 h. We chose to express Httex1(Q72) without tags such as RFP because we (27) and others (28–30) have observed that such tags alter the aggregation properties of mHttex1. Probing these cells with HD8-1 SP10-fluorescein (Fig. 6A) labels numerous intracellular round puncta, some of which are localized to the cytoplasm (arrows) and some in the nucleus (arrowheads). These structures were also labeled by the PHP1 antibody which binds within the PRD domain (epitope = QAQPLLPQP (31)) and reacts strongly with aggregated Httex1 (31) (Fig. 6B). The merged image (Fig. 6C) shows that PHP1 labels the outer shell of the puncta whereas HD8-1 SP10 fluorescence appears more uniform throughout, indicating that the peptide has access to the interior of the aggregates, perhaps due to its smaller size, while the antibody is more restricted to the puncta edge. Further, PHP1 labels the intranuclear aggregates poorly compared to cytoplasmic aggregates (Fig. 6B arrowheads) while HD8-1 SP10-fluorescein appears to label both strongly. Additionally, PHP1 and HD8-1 SP10 co-localization was also observed in peri-nuclear mesh like networks. However, some cells with diffuse PHP1 reactivity did not show HD8-1 SP10 fluorescence. These results validate that the peptide ligands bind Httex1 aggregates in cells, with more uniform staining than the well-established huntingtin directed PHP1 antibody. Further, the results indicate that the structural features recognized by the Httex1 binding peptides on *in vitro* made protofibrils are also present within misfolded Httex1 expressed in HEK293T cells and that these features are present throughout the puncta.

**Fig. 6.**
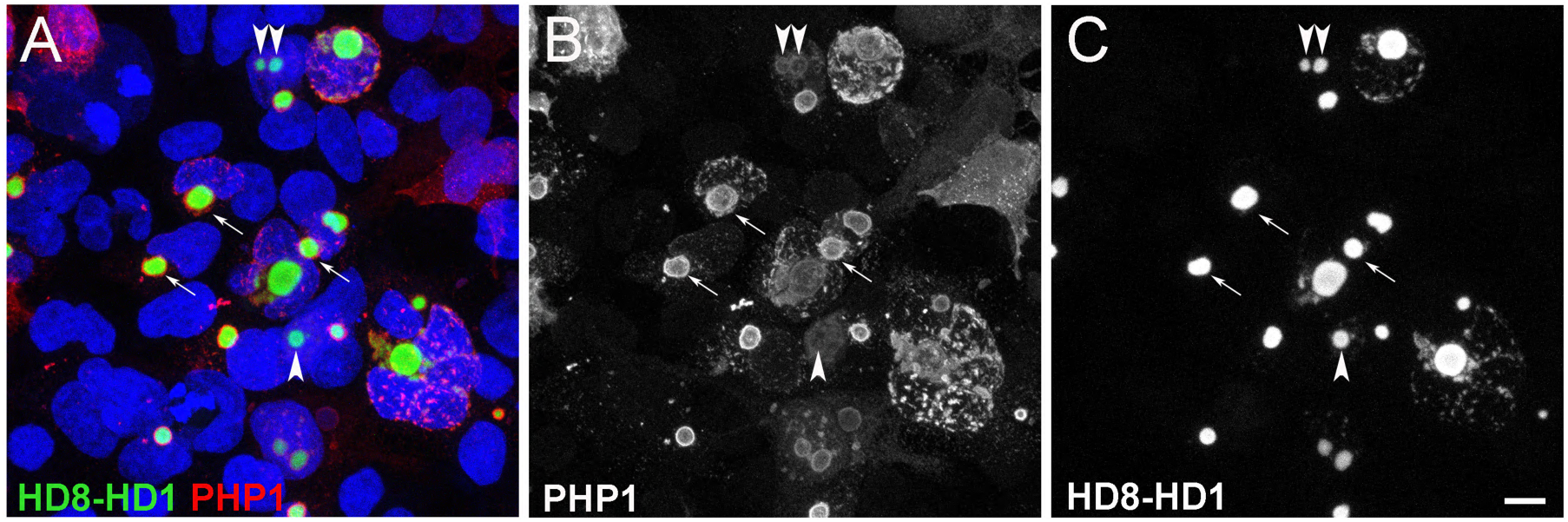
HD8-1 SP10 fluorescein and PHP1 antibody staining of Httex1Q72 aggregates expressed in HEK293T cells. *(A)* Merged image of HEK293T cells transfected with Httex1Q72 stained with HD8-1 SP10 fluorescein (green), PHP1 antibody (red), and nuclei stained with DAPI (blue). *(B)* PHP1 staining (white) and *(C)* HD8-1 SP10 fluorescein staining (white) of the same field. Arrows indicate cytoplasmic aggregates and arrowheads indicate intranuclear aggregates. Scale bar = 5 µm.

### GFP-HD8-1 SP10 Expression Reduces the Number of Httex1(Q72) Puncta Formed in Cells

The HD8-1 SP10 heterodimer strongly inhibited Httex1 aggregation *in vitro*. To see whether this ability is also present in a cellular context, we co-transfected HEK293T cells with Httex1(Q72) and GFP control (Fig. 7A) or with GFP-HD8-1 SP10 expression constructs (Fig. 7B). The number of GFP fluorescent cells showed similar transfection efficiencies in both conditions. Cells were fixed after 24 hours and incubated with PHP1 (Fig. 7, panels 2 and 5, magenta) and HD8-1-SP10-Alexa 647 to detect misfolded Httex1(Q72) (Fig. 7, panels 3 and 6, gray scale). In the wide field view, numerous PHP1 and HD8-1-SP10-alexa 647 positive small aggregates can be seen in the control GFP co-transfected condition (20X; Fig. 7 A panels 1-3). The magnified image (63X, Fig. 7A panels 4-6) illustrates the high degree of co-localization of PHP1 and HD8-1-SP10-alexa 647 signals in the round Httex1 puncta but not in cells with diffuse PHP1 labeling.

**Figure 7.**
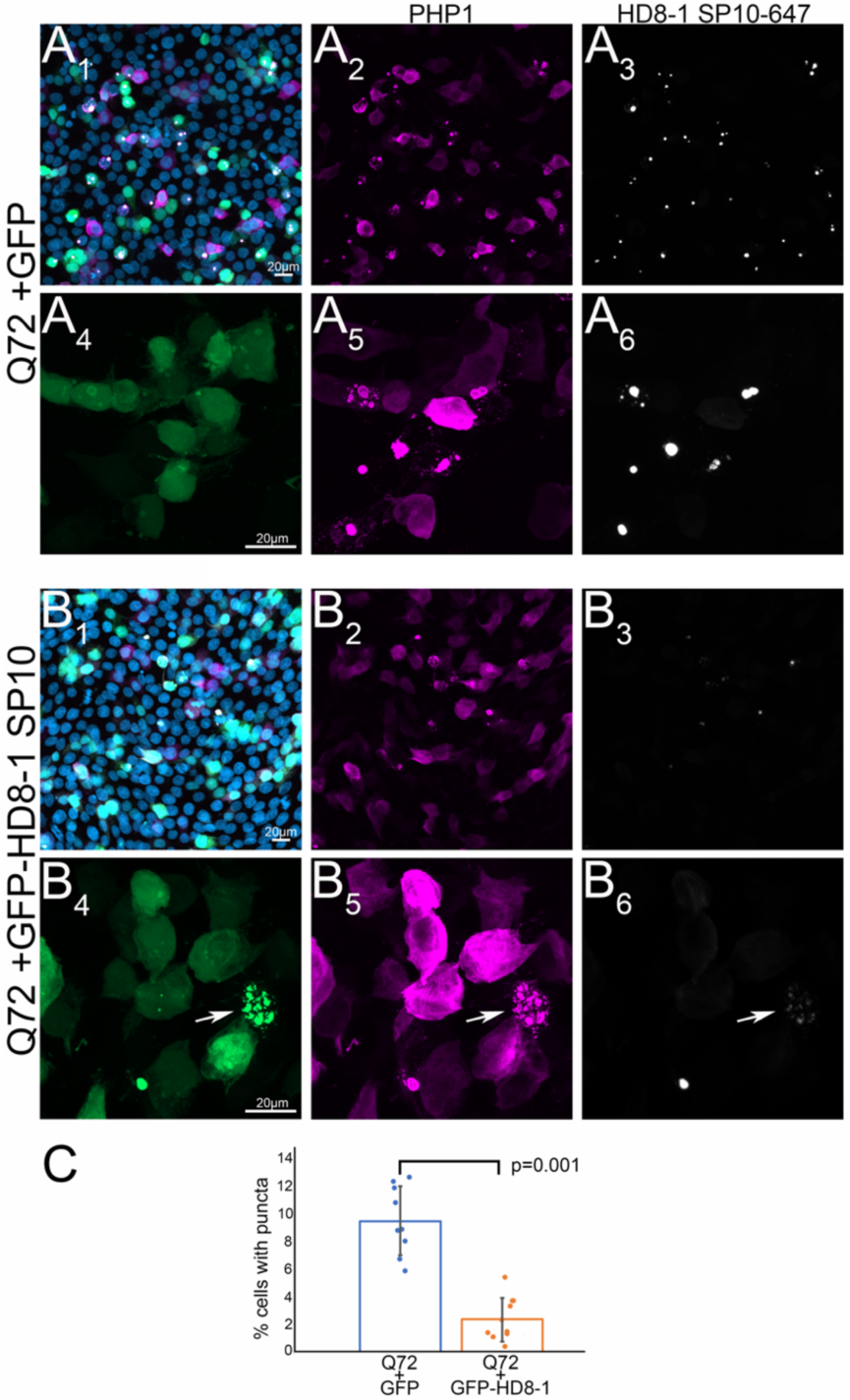
Coexpression of GFP-HD8-HD1 reduces the number of Httex1Q72 puncta formed in cells. Httex1Q72 was coexpressed in HEK293 cells with control reporter mEGFP (A_1-6_) or with mEGFP-HD8-HD1 SP10 (B_1-6_), a fusion protein with HD8-1 SP10 attached to the C-terminus of mEGFP. Httex1Q72 was visualized using PHP1 antibody (magenta) and HD8-1 SP10-Alexa 647 peptide (white) and nuclei were visualized with DAPI (blue) and panels A_1-3_ and B_1-3_ are low magnification (20X) and A_4-6_ and B_4-6_ are high magnification (63X) images across channels from the same fields. (A) Panel A_1_ is the merged image plus DAPI (blue) of panel A_2_ PHP1 (magenta) and panel A_3_ HD8-1 SP10-Alexa 647 (white). Panel A_4_ shows mEGFP indicating cells that were transfected and panels A_5_ and A_6_ show high magnification of fields in A_2_ and A_3_ respectively. (B) Panel B1 is the merged image plus DAPI (blue) of panel B2 PHP1 (magenta) and B3 HD8-1 SP10-Alexa 647 (white). Panel B_4_ shows mEGFP indicating cells that were transfected and panels B_5_ and B_6_ show high magnification of fields in B_2_ and B_3_ respectively. Arrows (B_4-6_) point to fragmented Httex1Q72 aggregates that colocalize with mEGFP HD8-HD1 SP10 (B_4_), PHP1 (B_5_) and HD8-1 SP10-Alexa 647 (B_6_). (C) Quantification of cells that contain Httex1Q72 puncta in cells expressing the mEGFP control or mEGFP-HD8-HD1 SP10. Cell number, based on DAPI stain, was quantified in 10 fields using ImageJ. The number and percent of cells with puncta was calculated and plotted for each group (see Materials and Methods) and compared using unpaired *t*-test. (Scale bar = 20 µM)

Interestingly, the size and number of PHP1 and HD8-1 SP10-alexa 647-positive round puncta were noticeably reduced when GFP-HD8-1 SP10 was co-transfected with Httex1(Q72) (Fig. 7B). Cells expressing GFP-HD8-1 SP10 also gave rise to more cells with diffuse PHP1 signal (Fig. 7B, panels 2 and 5). When present, aggregates appeared fragmented (Fig. 7B, panels 4-6; arrow), and smaller (c.f. panel A3 vs. B6). In addition to reducing puncta formation, GFP-HD8-1 SP10 shows good colocalization with PHP1-positive and HD8-1 SP10-alexa 647-positive Httex1 puncta that remained (Fig. 7 B4-6). Thus, endogenously expressed GFP-HD8-1 SP10 binds misfolded Httex1(Q72) but did not saturate all binding sites as evidenced by labeling by the exogenously added peptide. To quantify the aggregate-lowering effect, we determined the percent of cells containing HD8-1 SP10-alexa 647 positive Httex1 puncta (Fig. 7C). The results show a highly significant reduction in the number of puncta in cells co-transfected with GFP-HD8-1 SP10 (P=0.001, unpaired student *t-*test), corroborating the aggregation inhibition seen *in vitro*.

### HD8-1 SP10-fluorescein stains aggregated huntingtin in the R6/1 mouse model

Given that HD8-1 SP10 efficiently stains Httex1(Q72) transfected in cells, we next explored whether the HD8-1 SP10 peptide could be used to identify huntingtin aggregates in a mouse model system. We chose the R6/1 mouse that carries a human genomic fragment containing promoter sequences and exon 1 with ∼115 CAG expansion (8). The mutant Httex1 is ubiquitously expressed, and the mouse has a progressive neurological phenotype that mimics many features in Huntington’s disease, with symptoms onset at 15-21 weeks and death by ∼8 months. We looked at the neural retina because it does not accumulate lipofuscin granules which are abundant in aged brain and therefore simplifying fluorescent microscopy, and prior work had demonstrated the R6/1 mouse retinae exhibit huntingtin aggregates (32–34). Retinal cross sections were taken from 34-week-old R6/1 mice and control littermates, and stained with HD8-1 SP10-fluorescein, PHP1, and DAPI to visualize nuclei in the different retinal layers (Figure 8). Tissue from control littermate showed straight linear boundaries between retinal layers and no signal from HD8-1 SP10-fluorescein (Fig. 8A, panel 1), whereas PHP1, a mouse monoclonal antibody, labeled only retinal vessels due to the immunoglobulins present within (Fig. 8A, panel 2, magenta). In contrast to control retina, the R6/1 retina is often seen to contain folds, or retinal rosettes, due to disruption of polarity complexes at the apical junction of the outer nuclear (ONL) containing photoreceptor cell nuclei (34). In these retinal sections, HD8-1 SP10-fluorescein staining showed bright, punctate staining in all retinal layers (Fig. 8, B-D, yellow). At the ONL, most nuclei belong to rod cells (97%) with centrally located heterochromatin (35). Overlap of HD8-1 SP10-fluorescein and PHP1 signals can be seen in some cells (Fig. 8C, insets), although in most instances their signals do not overlap. A similar pattern can be seen at the inner nuclear layer (INL) that contains horizontal, bipolar and amacrine cells (Fig. 8C). At the ganglion cell layer (GC), intranuclear inclusions are labeled by both HD8-1 SP10-fluorescein and PHP1 (Fig. 8 D). HD8-1 SP10-fluorescein also labeled small and abundant cytoplasmic inclusions which were largely PHP1-negative. The absence of PHP1 signal may reflect the loss of epitope due to proteolysis. These data indicate that the structural features on the polyQ domain of the protofibrils that are binding sites for HD8-1 SP10 are present in the Httex1 aggregates in a model of human disease.

**Fig. 8.**
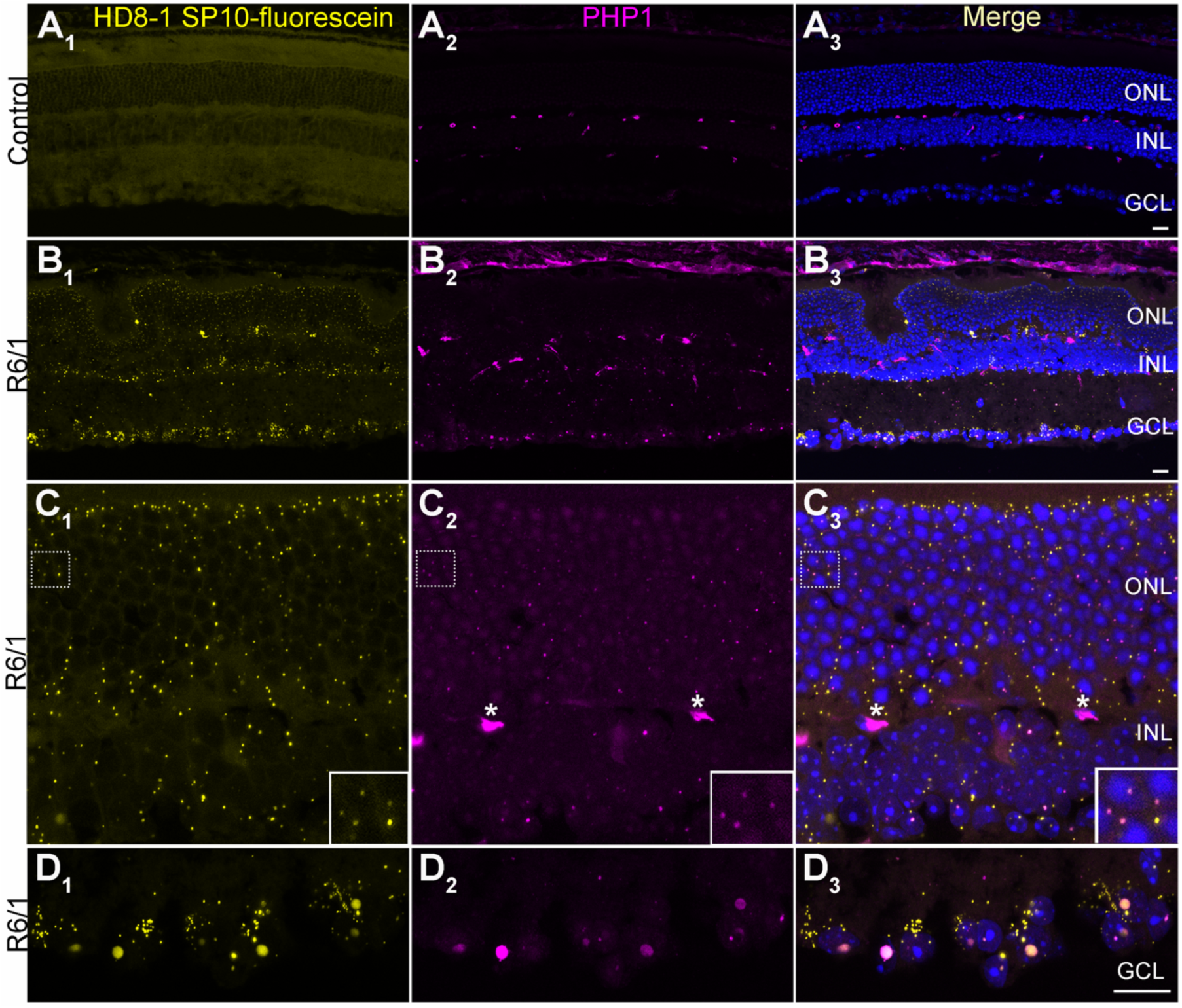
HD8-1 SP10 fluorescein staining of aggregated huntingtin in the retinae of the R6/1 mouse model. Retinal tissue sections from 34 week-old *(A)* age-matched control non-transgenic littermates (NT) and *(B-D)* R6/1 mice were stained with HD8-1 SP10 fluorescein (yellow; A_1_-D_1_), PHP1 (magenta; A_2_-D_2_), and with DAPI (blue, merged; A_3_-D_3_) as indicated. The retinal outer nuclear layer (ONL), inner nuclear layer (INL), and ganglion cell layer (GCL) are indicated. (A) No huntingtin-specific staining is observed for HD8-1 SP10 fluorescein (A_1,3_) or PHP1 (A_2,3_) in the control NT mice in any of the retinal layers. Non-specific labeling of retinal vessels is seen for the mouse monoclonal PHP1 antibody (A_2_,_3_; magenta). (B) R6/1 mouse retinae stained with HD8-1 SP10 fluorescein (B_1,3_) PHP1 (B_2,3_) and merged stained with DAPI (B_3_) at low resolution (scale bar = x microns). *(C)* Medium resolution imaging of the R6/1 ONL and INL retinal layers stained with HD8-1 SP10 fluorescein (C_1,3_) PHP1 (C_2,3_) and merged stained with DAPI (C_3_). Asterisks (*) indicate PHP staining of retinal vessels and the high resolution inset (boxed) expands the dashed area. (D) High resolution imaging the GCL layer stained with HD8-1 SP10 fluorescein (D_1,3_) PHP1 (D_2,3_) and merged stained with DAPI (D_3_) (scale bar = 20 µm). The high resolution inset (boxed) expands the dashed area.

## Discussion

### Selection Identifies Two Distinct Protofibril Binding Peptide Motifs

The mRNA display selection resulted in two major types of Httex1 protofibril binding peptides, the HD1 family (W-rich) and the HD8 family (Q and M rich) (Fig. 2A,B). Alignments of sequences within each family along with alanine scanning data identify key residues involved in protofibril recognition (Fig. 2 C,D) and define the regiospecificity of peptide protofibril recognition. Both peptides recognize Httex1 protofibrils by binding to structural epitopes only present in the misfolded polyQ region (Fig. 3A,B; Fig. S4). This distinguishes them from commonly used monoclonal antibodies, which typically detect Httex1 aggregates via epitopes in the proline-rich domain (PRD) - a region that does not undergo substantial structural rearrangement upon aggregation (13, 36). For example, antibodies such as PHP1 (31) and MW8 (37) bind to fibrils by interacting with the PRD, whereas polyQ-specific antibodies like MW1 (38) target monomeric Httex1 and fail to recognize the aggregated polyQ core of fibrils (38). Therefore, HD1, HD8, and HD8-1 SP10 (and related peptides) fulfill a critical need by directly detecting the formation of misfolded polyQ structures and specifically recognizing the toxic, protofibrillar form of Httex1 (13). We demonstrate the ability of these peptides to bind misfolded polyQ in cell culture and tissue from the R6/1 HD mouse model. The high specificity is evidenced by their overlapping signal with PHP1. Notably, HD8-1 SP10-fluorescein labeled more structures than PHP1, particularly in tissues (Fig. 8). This difference may be due to the smaller size of the peptide allowing for better access to the epitope. Alternatively, the epitope for PHP1 within the exposed PRD bristles may be lost due to proteolysis whereas the polyQ core should be more resistant.

### Peptide-Protofibril Recognition

mRNA display selection defines the protofibril surfaces compatible with high affinity peptide binding. The data argue that HD1 and HD8 bind to two different sites on the protofibril and that both sites are contained within the misfolded polyQ domain. First, the HD1 and HD8 sequences show essentially no sequence homology and very different sensitivities to alanine mutations. In HD1, tryptophan to alanine mutations eliminate more than 90% of the binding, while in HD8 glutamine to alanine mutations show the biggest impact. Second, the two peptides exhibit markedly different stoichiometries of binding to Httex1 protofibrils: HD8 binds nearly at a 1:1 ratio with Httex1, whereas HD1 displays a maximal stoichiometry of approximately 1:3 (Fig. 3). Both peptides are specific for protofibrils vs. the monomer or fully bundled fibrils (Fig. 1C; Fig. S5) and EPR data show that both HD1 and HD8 bind specifically to the misfolded poly Q region and that they bind protofibrils that lacking N17 (Httex1ΔN17) or the PRD (Httex1ΔPRD) (Fig. S4).

The peptide ligands also give insights into both Httex1 protofibril structure and rules for protofibril targeting. First, the observation that homodimers (HD1-1, HD8-8) and heterodimers (HD8-1,HD1-8) bind with much higher affinity than either monomer indicates there are multiple HD1 and HD8 sites within each protofibril and that these sites are relatively close in space. HD8 sites appear to be both more abundant and closer in space, given that a short linker (5 aa) gives optimal HD8-8 binding. On the other hand, there are about three times less HD1 sites and require a longer linker (10 aa) for optimal dimer binding (HD1-1, HD8-1, and HD1-8). Further, there appear to be different classes of HD1 sites, high (K_1_ = 210 nM; stoichiometry 1:50), medium (K_2_; stoichiometry 1:16) and low-affinity (K_3_ = 120 uM; stoichiometry 1:3.1). This presents a model where HD8-1 and HD1-8 dimers likely bind a high affinity HD1 site and a nearby HD8 site simultaneously. The other general feature we observe is a strong bias against positively charged residues (more than 10-fold over the selection) in both the HD1 and HD8 families (Fig. S3). The observed charge bias was somewhat unexpected, given that the polyQ region (the binding site for the HD peptides) carries no net charge. All charged residues in Httex1 are confined either to the extreme C-terminus of the PRD or to the N17 region. Due to the extended polyproline II helical conformation of the PRD, the C-terminal residues are positioned more than 10 nm away from the fibril core (36), likely rendering their charges functionally irrelevant for binding. In contrast, the N17 region lies in close proximity to the polyQ. Along with the N-terminal amine, it contains three lysines and two glutamates, resulting in an overall positive charge. Whether this net positive charge of N17 contributes to the observed charge bias remains to be determined.

HD1 sequences are most consistent with the peptide adopting a polyproline type II (PPII) left-handed helix. Proline and W residues occur at regular intervals (i, i + 3) and prolines strongly destabilize both beta sheets (39) and alpha helices (18). Searching for human homologs of HD1 identifies the thrombospondin type-1 motif (TSP-1) in AdamTS1. There, tryptophan residues adopt a polyproline type II helix and recognize two strands of a beta sheet by interdigitating between the R-, K-, and Q-rich sidechains (Fig. S8 A). This structure suggests that the repeating tryptophan residues in HD1 could interact similarly with glutamine beta sheet sidechains in Httex1 protofibrils. HD8-protofibril binding appears most compatible with the Q/M residues adopting a beta sheet configuration, similar to the interactions proposed by Perutz et al., (40) and seen in the Orb-2 structure (19). While this study primarily focused on peptide binders, the sequences responsible for binding could potentially be applicable to cellular proteins and help identify naturally occurring aggregate binders. For example, the fibril binding of peptides with Q10, Q15, and Q20 is consistent with the known incorporation of polyQ-containing, otherwise soluble proteins into fibrils (41–43). Similarly, HD1 and HD8 sequences might help to identify protofibril-binding proteins.

### Inhibition of Httex1(Q46) Misfolding

The HD1 and HD8-1 SP10 peptides are potent inhibitors of Httex1 misfolding, while HD8 has a much smaller inhibitory effect (Fig. 5 A-C). These observations argue that HD1 binding blocks the protofibril sites where fibril growth occurs. Further, increasing the affinity of HD1 by creating an HD8-1 dimer results in more potent misfolding inhibition. Indeed, HD8-1 SP10 represents the most potent Httex1 misfolding inhibitor described to date. Aggregation inhibition by HD1 occurs under conditions where only around 10% of the HD1 protofibril sites (high and medium affinity sites) are completely or partially occupied. Potent inhibition has also been observed for low stoichiometry small molecule protofibril binders (44). Together, these observations could be reconciled if high (K_1_), medium (K_2_) and low affinity (K_3_) HD1 protofibril sites correspond to fast, medium, and slow protofibril sites for Httex1Q46 aggregation and fibril growth. Overall, these results argue that optimizing protofibril binding provides a route to create potent misfolding inhibitors.

### HD8-1 SP10 binds Httex1 aggregates with high specificity and inhibits aggregate formation in cell culture model of HD

Previous ultrastructural studies on mHttex1 and polyQ aggregates show that they exhibit conformational heterogeneity (45). It is therefore important to see whether the cross-beta sheet structure of the polyQ region that form the HD8-1 binding sites are similar in protofibrils made in vitro and those made within HEK293 cells. This mammalian cell model system has been extensively used to investigate mechanisms of inclusion formation and pharmacologic agents that modulate this process (15, 46–48). We found that fluorescently labeled HD8-1 SP10 binds spherical inclusions in the nucleus and the cytoplasm with high specificity with little background fluorescence from non-transfected cells (Fig. 6). These round puncta were also recognized by PHP1, a validated antibody that prefers aggregated huntingtin (32). Under our staining conditions, PHP1 labels the shell of the puncta whereas HD8-1 SP10 labeling was more uniform throughout. It may be of relevance that ultrastructural studies of Httex1 inclusions formed in HEK293 cells reveal a distinctive core and shell morphology and are composed of highly organized fibrils (30). The staining pattern we observed with HD8-1 SP10 and PHP1 may reflect differential access of these binders to the shell and interior of the aggregates, with HD8-1 SP10 being substantially smaller in size than a mouse antibody. Alternatively, they may reflect different concentrations of binding targets at these locations. These results indicate that the structural elements in the *in vitro* formed protofibrils are present in the Httex1(Q72) round intranuclear and cytoplasmic inclusions.

HD8-1 SP10 potently inhibited Httex1(Q46) aggregation in vitro (Fig. 5). We tested its ability to inhibit aggregation in the cellular context by co-expressing GFP-HD8-1 SP10 and Httex1(Q72) in HEK293 cells. GFP was appended to the amino-terminus of HD8-1 SP10 to facilitate visualization. Indeed, we observed a significant and dramatic lowering of puncta (Fig. 7). Notably, GFP signal was observed in those cells with aggregates, indicating target engagement by HD8-1 SP10 despite the relatively large GFP (27 kD) being fused to the small peptide (3.3 kD). It is possible that the presence of GFP may have contributed to the efficacy of aggregate lowering due to steric interference with Httex1(Q72) fibril growth.

### Misfolded PolyQ as a Biomarker for mutant huntingtin

Formation of Httex1 aggregates in HEK293 cells takes hours whereas in animal models and humans aggregate formation takes months and years (49, 50) and may display structurally distinct variants. We investigated whether the binding targets of HD8-1 SP10 is present in the R6/1 mouse (8), a widely used HD animal model, at an age when aggregates are detectable with PHP1. Indeed, HD8-1 SP10-fluorescein labeled abundant cellular inclusions in R6/1 tissue but not in age-matched littermate controls (Fig. 8). The staining pattern appears to be highly specific, with the large intranuclear aggregates in the staining pattern that partially overlapped with PHP1 (Fig. 8). Notably, in the ganglion cells, HD8-1 SP10 brightly labeled numerous small cytoplasmic puncta whereas signal from the intranuclear aggregates was relatively weaker. In contrast, PHP1 signal was more noticeable in the intranuclear inclusions compared to those in the cytoplasm.

Overall, this work provides new molecular tools to recognize misfolded polyQ as well as a route to inhibit mutant huntingtin aggregation *in vitro* and in *vivo*.

## Materials and Methods

### Expression and purification of Httex1 fusion proteins and mutants

Httex1(Q46) was expressed recombinantly in *E. coli* BL21 DE3 as a thioredoxin fusion protein (thioredoxin-Httex1Q46-His_6_) cleaved from the fusion tag and purified using previously published protocols (11). A cysteine mutant (Httex1-C111) was expressed for labeling with spin label or biotin. Httex1 mutants with deleted N17 or PRD regions (ΔN17-Htt46Q (thioredoxin-KKQ46PRD-His_6_) and Htt25Q-ΔPRD (thioredoxin-N17-25Q-KK-His_6_)) were used in truncation experiments. Briefly, Httex1 was purified using ion-exchange chromatography, cleaved with enterokinase EKMax (Cat# E18002, ThermoFisher Scientific) to remove the thioredoxin tag, and fractionated using a Phenomenex C4 column (A=99.9% water 0.1% trifluoroacetic acid; B = 90% acetonitrile, 9.9% water and 0.1% TFA). The protein was then lyophilized and stored under vacuum until further use. Protofibrils were formed as described previously (13). Fibrillation was initiated by adding ice-chilled TBS buffer (20 mM Tris, 150 mM NaCl, pH 7.4) to the Httex1 protein film to give a final protein concentration of 15 µM. Fibrillation was carried out overnight at 4^°^ C, after which the fibrils were harvested by ultracentrifugation, resuspended in 0.025% trifluoroacetic acid in water (TFA-Water), and sonicated using a tip sonicator (Microson ultrasonic cell disruptor, XL2000) 3-5 times on ice for 10 seconds to obtain protofibrils. (13).

### Preparation of biotin-containing protofibrils and immobilized Httex1

Httex1 protofibrils were constructed by combining two versions of Httex1 during fibril formation, Httex1(Q46) (WT) and Httex1(Q46)-C111-biotin. Httex1(Q46)-C111-biotin is identical to Httex1(Q46) with one modification, a H111C mutation which was subsequently biotinylated. The fusion protein was incubated with 20-fold molar excess of EZ-Link Maleimide-PEG11-Biotin (Cat # 21911, ThermoFisher Scientific) in TBS (20 mM Tris, 150 mM NaCl, pH 7.4) at room temperature for two hours to biotinylate the cysteine moieties. Biotinylated fusion protein was then purified using ion-exchange chromatography, cleaved with enterokinase EKMax (Cat # E18002, ThermoFisher Scientific) to remove thioredoxin fusion partner and fractionated over C4 reversed-phase column to obtain purified biotinylated Httex1 (Httex1-C111-Biotin). The reversed-phase purified Httex1-C111-Biotin was lyophilized and stored in vacuum desiccator until further use. To create biotinylated protofibrils, unmodified Httex1(Q46) was combined with Httex1(Q46)-C111-biotin (20:1 ratio), fibrils were formed, sonicated, and quantified using published methods (13).

### Preparation of Immobilized Httex1 Protofibrils for mRNA Display Selection and Binding analysis

Immobilized Httex1 protofibrils were generated by dissolving Httex1(Q46)-C111-biotin protofibrils in immobilization buffer (50 mM NaOAc, pH 4.0, 150 mM NaCl, and 0.1% Tween-20) and streptavidin agarose beads (Thermo Fisher Scientific cat# 20349) or Streptavidin Ultralink resin (Thermo Fisher Scientific cat# 53113). Concentrations indicate the concentration of Httex1 monomer immobilized on the bead as protofibril.

### Preparation of Immobilized Httex1 Monomer

20 uL streptavidin agarose beads were washed 3X with immobilization buffer (50 mM NaOAc, pH 4.0, 150m NaCl, 0.1% Tween20), mixed with 420 pmol of Httex1 monomer, and incubated at 4°C for 3 hours. The beads were then washed 3X with selection buffer (20 M Tris, pH7.6, 150 mM NaCl, 0.1% Tween20, 1 mg/mL BSA and 50 ug/mL tRNA), blocked in selection buffer plus 20 uM biotin at 4C for 1 hour and washed.

### Preparation of Immobilized α-synuclein Protofibrils

Biotinylated α-synuclein fibrils, wild-type α-synuclein and a single cysteine mutant with a cysteine residue at position 136 were expressed and purified as previously described (51). The C136 α-synuclein mutant was biotinylated as described for Httex1 above. Biotinylated α-synuclein fibrils were generated by mixing biotin labeled, α-synuclein-136-biotin with wild type alpha synuclein in a 1:20 ratio. Fibrils were gown at a total protein concentration of 200µM in TBS buffer (20 mM Tris, 150 mM NaCl, pH 7.4) for 7 to 10 days at 37degree using stir bars as described (51).

### mRNA display Selection

A naïve mRNA display library (MX_9_-GSGTSGSS) with approximately 1 x 10^12^ independent sequences was generated using rabbit reticulocyte lysate and a mRNA display selection was performed essentially as described previously (52) using immobilized Httex1 protofibrils as the target. Eight rounds of selection were performed. In rounds one to four, 280 pmol (monomer) of Httex1 protofibrils were immobilized on 30 uL of resin. In rounds five to eight, the target was reduced to 90 pmol (monomer) protofibrils and 10 μL resin. In each round, the cDNA/mRNA-peptide fusion library was added to immobilized Httex1 unbundled fibrils in 1 mL of selection buffer (20 mM Tris-HCl, pH 7.6, 150 mM NaCl, 1 mg/mL BSA, 50 μg/mL tRNA, 0.1% Tween-20, and 20 μM Biotin) with rotation at 4 °C for one hour (rounds one to five) or 37°C (round six to eight). Target-specific binding was monitored using radioactive pulldown assays. Radioactive mRNA-peptide fusions were generated using *in vitro* translation with [^35^S] Met (53, 54). Radiolabeled peptide-fusions were purified with oligo dT, treated with RNaseA, and added to Httex1 immobilized unbundled fibrils (either NutrAvidin® agarose or Streptavidin UltraLink®), incubated in 1 mL of selection buffer at 4 °C for 1 hour with rotation. The resin was washed three times with selection buffer, and the binding of peptides was determined by scintillation counting of radioactive counts. Net binding was observed after two rounds of enrichment (pool 2).

### Sequence Analysis and Clone Identification

cDNA from pools six, seven, and eight were sequenced manually and via high throughput sequencing to give protofibril binding peptides (HD1 – HD8). Approximately four million reads were obtained from pools six, seven, and eight (USC Molecular Genomics Sequencing Core), curated to remove stop codons and deletions, and the top 419 sequences in each pool were identified (≥ 500 matching reads). These were binned by sequence similarity using the HAMMOCK program (55) giving four major groups—two that were tryptophan rich and two containing multiple glutamines/methionines. The highest abundance members of each group were tested for binding to immobilized protofibrils (Sup. Fig. 1B) two peptides, HD1 (tryptophan rich) and HD8 (glutamine rich) showed the highest binding to the protofibril targets.

### Sequence Composition and Alignment

In the round 8 pool, approximately 200 sequences correspond to the W-Rich or Q-Rich groups. The amino acid composition in the round 8 pool was determined and compared to the starting library composition (the nine randomized positions in the MX_0_ library) and plotted as the log_10_ (pool 8 count/pool 0 count) such that no change in composition between pool 0 and pool 8 = 0 (Fig. 2). Each group was further subdivided by the position of either the W or Q residues, the sequences binned by the position of the W or Q residues and then visualized using the Weblogo online tool (https://weblogo.berkeley.edu/logo.cgi) (56, 57). W-rich sequences were subdivided into five groups: i) W3,W4,W7, ii) W4,W7, iii) W4, W7, W10, iv) W7, and v) W7, W10. The Q-rich peptides are more diverse and give at least 19 distinct groups based on the Q positions (e.g., “HD8-like” = Q3,Q10; Sup. Fig. 1). These 19 groups can be binned into four larger groups by the position of conserved Q or M residues: i) “HD-8 like” Q/M at positions 1, 3, 5, 8, and 10 (Fig. 2B), ii) Q/M at positions 1, 3, 5, and 7, iii) Q/M at positions 1, 4, 6, and 8, and 10 iv) Q/M at positions 1, 3, 5, 6, 8, 10.

### Binding Analysis by Radioactive Pulldown

Pool and individual clone binding was assayed by radioactive pulldown using RNase-treated peptide fusions as described (24). Individual peptides or selection pools were labeled with [^35^S]-methionine by *in vitro* translation using rabbit reticulocyte lysate. Radiolabeled samples were purified with oligo dT, eluted, and treated with RNase to give F30P DNA linker-peptide fusions (24). Binding assays were performed by incubating the labeled fusions with Httex1 protofibrils immobilized on resins (either NeutrAvidin® agarose or Streptavidin UltraLink®), washing 3X with selection buffer, and quantitating binding by scintillation counting.

### Alanine Scanning

Single alanine mutations were introduced for each of the first ten residues in HD1 and HD8 (21) by synthesizing each construct at the DNA level using oligonucleotides (IDT). To probe the N-terminal methionine, an additional alanine residue as added between the first and second positions

### Solid Phase Peptide Synthesis

All peptides were synthesized via FMOC solid phase chemistry on Rink amide resin using an automated synthesizer (Biotage Syro, USC Center for Peptide and Protein Engineering-USC-CPPE) or obtained commercially. HD1 (MDWWPMWPSL), HD8 (MWQMMNGMSQ), HD1-G (MDWWPMWPSLG), HD8-G (MWQMMNGMSQG), HD1GGEK (MDWWPMWPSLGGEK), (HD1-8 SP10 (MDWWPMWPSLGSGTSGSSGSMWQMMNGMSQ), HD8-1 SP10 (MWQMMNGMSQGSGTSGSSGSMDWWPMWPSL), and HD1-1 SP10 (MDWWPMWPSLGSGTSGSSGSMDWWPMWPSL) were synthesized on Rink Amide 4-methylbenzhydrylamine hydrochloride (MBHA) resin at the 10 or 25 umol mmol scale The automated synthesizer was provided with stock solutions of Fluorenylmethyloxycarbamate (Fmoc)-protected amino acids (0.5M in NMP), 4-methylpiperidine (40% v/v in NMP), 1-[Bis(dimethylamino)methylene]-1H-1,2,3-triazolo[4,5-b] pyridinium 3-oxide hexafluorophosphate (HATU) (0.49M in NMP), and N,N-diisopropylethylamine (DIEA) (2M in NMP), and NMP as system solvent. Before beginning synthesis, dry resin was swelled in 5 mL of N-methyl-2-pyrrolidone (NMP) at room temperature for 30 minutes with intermittent vortexing, then drained before proceeding with synthesis. For Fmoc deprotection, 400uL of 40% 4-methylpiperidine was added to the resin, reacted at 55 °C for 5 minutes, then drained. An additional 200uL of 40% 4-methylpiperidine and 200uL of NMP were added, reacted at 55 °C for 10 minutes, then drained, and the resin washed five times with 900uL NMP. Amino acid couplings were carried out by mixing 200uL of 0.5M Fmoc-protected amino acid (4 equivalents), 210uL of 0.5M HATU (4.2 equivalents), and 100uL of 2M DIEA (8 equivalents). Coupling reactions were heated for 15 minutes at 55 °C, then drained and the resin washed five times with 900uL NMP. Fmoc deprotection of the final residue was performed with 4-methylpiperidine as described above. After synthesis, the resin was washed three times with 5mL dichloromethane (DCM).

Crude peptide was cleaved from the resin and side chain protecting groups removed with 8 mL of cleavage cocktail [94% (v/v) Trifluoroacetic acid (TFA), 2.5% (v/v) 2-2’-(Ethylenedioxy)-diethanethiol (DODT), 2.5% (v/v) deionized water, 1% (v/v) Triisopropylsilane (TIS)] for 2.5 hours at room temperature. The crude peptide was then precipitated in 40mL of ice-cold methyl-tert-butyl-ether (MTBE), pelleted by centrifugation, and dried by evaporation (Biotage V-10 Touch) before dissolving in dimethyl sulfoxide (DMSO) for purification. Peptides were purified by HPLC/MS (InfinityLab 1260, Agilent) using a C-18 reverse phase column (Aeris 5um PEPTIDE XB-C18 100Å, 250 x 10.0 mm, Phenomenex) with gradient elution [Solvent A: 95% (v/v) H2O, 5% (v/v) CH3CN, 0.1% (v/v) formic acid; Solvent B: 90% (v/v) CH3CN, 10% (v/v) H2O, 0.1% (v/v) formic acid]. Fractions were analyzed by in-line electrospray ionization mass spectrometry, and confirmed fractions were pooled and dried by evaporation.

Peptides for EPR studies were synthesized with a GGC linker to enable coupling with thio-reactive spin label and coupled as previously described (11). Fluorescein-labeled HD8-1-SP10 (MWQMMNGMSQGSGTSGSSGSMDWWPMWPSLGGC-Fluor) was synthesized as above, purified, and coupled with fluorescein-maleimide (CAS # 75350-46-8; Anaspec catalog number AS-81405 or Thermo Scientific™ catalog number 62245). Peptide (1 eq) was combined with fluorescein-maleimide (5 eq) in 0.5mL of DMSO, then mixed with 0.5 mL of 100 mM HEPES-KOH, pH 7.5, containing 2 equivalents TCEP (tris(2-carboxyethyl)phosphine) to prevent disulfide formation. Reactions were covered from light, then left at room temperature for 2-5 hours. Labeled peptides were purified and analyzed by HPLC/MS as above. HD8-1-SP10-Alexa 647 was contained Alexa Fluor™ 647 linked through maleimide (cat.# A20347, Thermo Fisher). HD1 and HD8 for competition experiments were synthesized with a single C terminal G (HD1-G and HD8-G).

### Stoichiometry Determination and Protofibril Binding by EPR

To determine the binding stoichiometries of HD peptides with respect to protofibrils, the EPR spectra of spin labeled HD peptides were monitored in the presence or absence of protofibrils. Protofibrils were obtained as described above. HD peptides were labeled by appending a GGC sequence at the C-terminus of HD1 and HD8. Peptides were obtained by solid state peptide synthesis. Peptide powders were dissolved to a concentration of ∼6 mM in a 50% DMSO/water solution containing 5-fold molar excess of MTSL spin labeling reagent (1-oxyl-2,2,5,5 tetramethyl-D3-pyrroline-3-methyl) methanethiosulfonate (Toronto Research Chemicals). The reaction was quenched by running the reaction mixture on a Phenomenex C4 reverse phase column using a gradient generated from buffer A (99.9% water and 0.1 trifluoroacetic acid) and buffer B (90% acetonitrile, 9.9% water and 0.1% TFA). Labeled peptides were identified based on their characteristic EPR signals and lyophilized. EPR spectra of labeled peptides in solution were obtained in 20 mM Tris, 150 mM NaCl, pH 7.4. The spectra of the fully protofibril-bound states were obtained from a mixture of 10 μM peptide in the presence of saturating amounts of protofibrils (300 μM). Small amounts of residual monomeric peptides were subtracted. Binding stoichiometries were estimated by incubating varying amounts of peptide in the presence of 8 μM protofibrils and recording the respective EPR spectra. For each of the spectra, relative amounts of mobile and immobile spectral components were determined using double integration methods (22). All experiments were done in triplicate.

### Peptide Binding Isotherms to Httex1(Q46) Protofibrils

Binding isotherms for HD1, HD8, and HD8-SP10-HD1 were measured at room temperature by [^35^S]-Met radioactive pulldown assay of RNase treated fusions as described above, eliminating the washing steps, and determining the fraction bound by scintillation counting the beads and supernatant. Six different Httex1 protofibril concentrations were used for the pulldowns (0 nM, 250 nM, 500 nM, 1000 nM, and 4000 nM for HD1 and HD8; 0 nM, 10 nM, 50 nM, 100 nM, 200 nM, and 400 nM for HD8–SP10–HD1), where protofibril concentration indicates the molarity of the total immobilized Httex1. Httex1 protofibrils were immobilized on Streptavidin UltraLink® resin by adding 100 pmol Httex1 as protofibrils per ul of resin (for HD1 and HD8) and 50 pmol Httex1 as protofibrils per ul of resin (HD8-SP10-HD1). The appropriate amount of Httex1 bound resin was added to each tube and the total amount of resin was adjusted to 10 μL by adding blank resin. The radiolabeled peptide fusions were incubated with the resin at room temperature for 24 hours in 250 μL of selection buffer. After the incubation, beads and supernatant were separated using Spin-X columns, and radioactive counts of each sample were determined by scintillation counting. Binding constants were determined using GraphPad Prism.

### Steady State Surface Plasmon Resonance Analysis of HD1

SPR experiments were conducted using a Biacore T200 instrument (Cytiva), using a Series S SA chip (Cytiva), at the USC Nanobiophysics core. The running buffer was TBST (20 mM Tris-HCl pH 7.5, 150 mM NaCl, 0.1 % (v/v) Tween-20) containing 1mg/mL bovine serum albumin (BSA). Running buffer was prepared on the day of the experiment and filtered immediately prior to use. The HD1-GGEK variant (MDWWPMWPSLGGEK) was used to ensure the solubility of HD1 at high peptide concentrations in the absence of DMSO. HD1-GGEK concentrations were determined by absorbance at 280nm, (extinction coefficient = 17070 M^-1^cm^-1^). Immediately before use, a frozen stock solution of biotinylated Httex1 was thawed, diluted in TBST to approximately 0.005mg/mL (∼350nM as monomer), and immobilized on the chip by manual injection to a surface density of ∼1200 Response Units (RU). HD1-GGEK peptide was used as the analyte in concentrations ranging from 60 nM to 240 uM at 25°C in TBST Buffer (20 mM Tris, 150 mM NaCl, 0.1% v/v Tween 20, pH 7.4) plus 1mg/mL BSA. The instrument was operated in multi-cycle kinetics mode, with sufficient time between injections for full dissociation. Peptide samples at the indicated concentrations were injected at a flow rate of 50 uL/min, at 25 °C, in triplicate. Each peptide was injected for a contact time of 60 sec, followed by 180 sec of dissociation. Blank injections consisted of running buffer alone, and the average blank response was subtracted from all raw values for baseline correction. Data were analyzed using the Biacore Evaluation software, Microsoft Excel, and Graphpad Prism. Curve fitting was performed in GraphPad Prism and Kaleidagraph (Synergy Software).

### K_d_ and Scatchard Analysis

The response vs. concentration plot did not fit well with a single K_d_ model at low concentration values, so two and three site models were considered. Scatchard analysis (23, 25) using an amplitude of 53 RU (The m.w. of Httex1(Q46) is 13030 g/mol and HD1GGEK is 1718 g/mol, indicating a 1:3 ratio of peptide:Httex1 binding should give an amplitude of 53 RU) resulted in a highly curved plot, consistent with at least two distinct sites and two microscopic binding constants (K_1_ and K_3_) between the peptide and immobilized protofibrils. Estimates for the high affinity (K_1_) and low affinity (K_3_) microscopic binding constants were made by calculating the limiting slope at ν = 0 and ν = 1 (where the slope = -1/K), giving k_1_ = 210 nM and K_3_ = 120 uM and iterated to determine the fraction of each site (ν_1_ and ν_3_), as described by Schimmel and coworkers (25, 26). This two-site model did not fit the Scatchard plot at intermediate values of ν (∼0.2), so a three-site model was used by including an intermediate binding constant and fraction giving K_1_ = 210 nM and ν_1_ = 0.024, K_2_ = 16 uM and ν_2_ = 0.075, and K_3_ = 120 uM and ν_3_ = 0.90.

### Aggregation kinetics monitored by EPR

A Q35C mutant of Httex1(Q46) was spin labeled to give rise to the Httex1(Q46)-35R1 derivative as previously described (11). To start the aggregation, ice-chilled TBS buffer was added to a dried protein film of lyophilized Httex1(Q46)-35R1 to obtain a 10 μM solution of Httex1(Q46)-35R1. To determine the effect of HD peptides on aggregation kinetics, HD peptides were added to the protein solution at a final concentration of 20 μM immediately after dissolving the protein film in TBS. The aggregation kinetics were obtained by monitoring the EPR signal amplitude of the central line over time (11). Amplitudes for each aggregation reaction were normalized to their respective initial values.

### Co-localization of Httex1(Q72) aggregates with HD8-1 SP10 peptides and PHP1 antibody

HEK293T cells were split and seeded onto 24-well plates with poly-D-lysine coated coverslips and grown in DMEM supplemented with 10% FBS until 75% confluency. Httex1Q72 plasmid DNA (1 µg), mixed with Lipofectamine, were applied on cells, and left for 4h according to the Lipofectamine LTX DNA transfection reagent protocol (Invitrogen). Twenty-four hours after transfection, cells were rinsed with PBS, and then fixed with 3.7% PFA for 10 minutes at room temperature. The cells were rinsed with the PBS and incubated in 0.1%Triton X-100 for 15 min followed by 1% BSA/PBS for one hour at room temperature. The primary antibody PHP1 (1:200, MABN2490, Millipore Sigma) was diluted in 0.1% BSA/PBS and incubated for overnight at 4°C. The next day, the coverslips were washed with PBS, and incubated with anti-mouse -594 (1:400, Thermo Fisher Scientific Cat # A-21203), and peptide (HD8-1 SP10-fluorescein, final concentration 0.5 µM) in 0.1% BSA/PBS and incubated for 2 hours at room temperature. The cells then mounted on a glass with mounting media with DAPI and were imaged with Zeiss LSM 800 confocal microscope.

### Lowering of the number of Httex1Q72 puncta in HEK293T cells by GFP-HD8-1 SP10

Httex1(Q72) and GFP-HD8-1 SP10 were cloned into pcDNA 4.0/TO mammalian expression vector (Thermo Fisher Scientific). HEK293T cells were split and seeded onto 24-well plates with poly-D-lysine coated coverslips and grown to 75% confluency. Cells were then co-transfected with 250ng of Httex1(Q72) and 250ng of mEGFP or GFP-HD8-1 SP10 plasmid DNA, according to the Lipofectamine LTX DNA transfection reagent protocol (Invitrogen). After 24 hours, cells were washed and fixed with 3.7% formaldehyde for 15 minutes and blocked with 1% bovine serum albumin (BSA) for 1 hour at room temperature. Cells were incubated with mouse PHP1 primary antibody (1:500) in 0.1% BSA/PBS blocking buffer overnight at 4°C, followed by incubation with Alexa Fluor 594 donkey anti-mouse IgG (1:200, Invitrogen), HD8-1 SP10-Alexa 647 peptide (1 µM, Thermo Fisher) and DAPI (0.5 µg/mL) in the same blocking buffer for 1 hour at room temperature. Coverslips were mounted onto glass micro slides with Vectashield antifade mounting medium (Vector Laboratories) and imaged with Zeiss LSM 800 confocal microscope. ImageJ was used to count the number of cell nuclei, based on DAPI stain (405nm channel) across 10 fields. The number of Httex1(Q72) puncta was manually counted per field using the HD8-1 SP10-Alexa 647 stain (647nm channel). The percent of cells with puncta was calculated and plotted for each group and numbers were compared using unpaired student *t-*test.

### Retinal sections immunofluorescence

The R6/1 (JAX stock #006471) and non-transgenic (NT) control mice were euthanized at 34 weeks of age. Eyes were enucleated and the cornea removed. The eyecups were fixed with 4% formaldehyde in PBS for 1-2 hours on a shaker at room temperature then washed 3 times with PBS (10 min each). Tissue was cryoprotected by 1 hour incubation with 15% sucrose, and then with 30% sucrose for overnight at 4°C. On the next day, the lens was taken out, the eyecups were embedded with OCT, and rapidly frozen either in liquid nitrogen or dry ice/ethanol and stored in -80℃. Frozen sections (10 µm) were obtained using a cryostat (Leica CM3050S). Sections were stored at -80°C. Prior to incubation with antibody or peptides, retinal cryosections were air-dried. A blocking buffer (5% normal donkey serum, 0.3%Triton X-100 and 1% BSA in PBS) was applied for 1h at room temperature. The sections were then incubated with PHP1 (1:200 diluted in blocking buffer) overnight at 4℃. The next day, sections were washed (3x5 min with PBS), and incubated with 1 µM HD8-1-SP10-fluorescein diluted with blocking buffer and Alexa Fluor 594-conjugated anti-mouse antibody (1:200, Invitrogen) for 2 hours at room temperature, followed by three washes in PBS. The sections were coverslipped with mounting media containing DAPI (Vectashield, Vector Laboratories). All images were taken with LSM 800 (Zeiss) confocal microscope.

## Supporting information

Supporting information for Noridomi et al 2025

## Acknowledgments

Peptide synthesis, purification, and analysis was performed at the USC Center of Excellence in Peptide and Protein Engineering (USC-CPPE) (https://sites.usc.edu/cppe/).

SPR experiments were conducted at the USC Center of Excellence in Nanobiophysics (https://dornsife.usc.edu/nanobiophysicscore/)

Peptides were purified and analyzed at the Agilent Center Of Excellence In Biomolecular Characterization (https://dornsife.usc.edu/agilentcoe/) at USC.

High throughput sequencing was performed at the USC Molecular Genomics Core in the USC Norris Comprehensive Cancer Center (https://uscnorriscancer.usc.edu/molecular-genomics-core/).

Optical Imaging and Microscopy was performed at the Optical Imaging Facility (https://microscopy.usc.edu/) located in the USC Stem Cell Institute (USC Stem Cell; https://stemcell.keck.usc.edu/).

## Funding

This work was supported by funded by CHDI (grant # A-15631) to R.L., NIH (R01NS125769) to R.L., R.W.R., and J.C., NIH EY12155 (J.C.), and the National Eye Institute Core Grant P30EY029220 to the University of Southern California, Department of Ophthalmology.

## Notes

**Competing Interest Statement:** A patent application WO2022051673A1 has been filed.

### Competing Interest Statement

A patent application WO2022051673A1 has been filed.

## References

1. G. P. Bates et al., Huntington disease. Nat Rev Dis Primers 1, 15005 (2015).

2. D. R. Langbehn, M. R. Hayden, J. S. Paulsen, P.-H. D. I. o. t. H. S. G. and the, CAG-repeat length and the age of onset in Huntington disease (HD): a review and validation study of statistical approaches. Am J Med Genet B Neuropsychiatr Genet 153B, 397–408 (2010).

3. P. McColgan, S. J. Tabrizi, Huntington’s disease: a clinical review. Eur J Neurol 25, 24–34 (2018).

4. E. Scherzinger et al., Huntingtin-encoded polyglutamine expansions form amyloid-like protein aggregates in vitro and in vivo. Cell 90, 549–558 (1997).

5. K. Sathasivam et al., Aberrant splicing of HTT generates the pathogenic exon 1 protein in Huntington disease. Proc Natl Acad Sci U S A 110, 2366–2370 (2013).

6. M. DiFiglia et al., Aggregation of huntingtin in neuronal intranuclear inclusions and dystrophic neurites in brain. Science 277, 1990–1993 (1997).

7. A. Lunkes et al., Proteases acting on mutant huntingtin generate cleaved products that differentially build up cytoplasmic and nuclear inclusions. Mol Cell 10, 259–269 (2002).

8. L. Mangiarini et al., Exon 1 of the HD Gene with an Expanded CAG Repeat Is Sufficient to Cause a Progressive Neurological Phenotype in Transgenic Mice. Cell 87, 493–506 (1996).

9. R. Wetzel, Exploding the Repeat Length Paradigm while Exploring Amyloid Toxicity in Huntington’s Disease. Acc Chem Res 53, 2347–2357 (2020).

10. M. Jayaraman et al., Slow amyloid nucleation via alpha-helix-rich oligomeric intermediates in short polyglutamine-containing huntingtin fragments. J Mol Biol 415, 881–899 (2012).

11. N. K. Pandey et al., The 17-residue-long N terminus in huntingtin controls stepwise aggregation in solution and on membranes via different mechanisms. J Biol Chem 293, 2597–2605 (2018).

12. K. Shen et al., Control of the structural landscape and neuronal proteotoxicity of mutant Huntingtin by domains flanking the polyQ tract. Elife 5 (2016).

13. J. Mario Isas et al., Huntingtin fibrils with different toxicity, structure, and seeding potential can be interconverted. Nat Commun 12, 4272 (2021).

14. M. Arrasate, S. Mitra, E. S. Schweitzer, M. R. Segal, S. Finkbeiner, Inclusion body formation reduces levels of mutant huntingtin and the risk of neuronal death. Nature 431, 805–810 (2004).

15. Y. E. Kim et al., Soluble Oligomers of PolyQ-Expanded Huntingtin Target a Multiplicity of Key Cellular Factors. Mol Cell 63, 951–964 (2016).

16. G. Kamalinia, B. J. Grindel, T. T. Takahashi, S. W. Millward, R. W. Roberts, Directing evolution of novel ligands by mRNA display. Chem Soc Rev 50, 9055–9103 (2021).

17. Y. Nagai et al., Inhibition of polyglutamine protein aggregation and cell death by novel peptides identified by phage display screening. J Biol Chem 275, 10437–10442 (2000).

18. A. Chakrabartty, T. Kortemme, R. L. Baldwin, Helix propensities of the amino acids measured in alanine-based peptides without helix-stabilizing side-chain interactions. Protein Sci 3, 843–852 (1994).

19. R. Hervas et al., Cryo-EM structure of a neuronal functional amyloid implicated in memory persistence in Drosophila. Science 367, 1230–1234 (2020).

20. J. E. Barrick, R. W. Roberts, Sequence analysis of an artificial family of RNA-binding peptides. Protein Science 11, 2688–2696 (2002).

21. B. C. Cunningham, J. A. Wells, High-Resolution Epitope Mapping of hGH-Receptor Interactions by Alanine-Scanning Mutagenesis. Science 244, 1081–1085 (1989).

22. H. Moaven et al., Visual arrestin interaction with clathrin adaptor AP-2 regulates photoreceptor survival in the vertebrate retina. Proc Natl Acad Sci U S A 110, 9463–9468 (2013).

23. G. Scatchard, The Attractions of Proteins for Small Molecules and Ions. Annals of the New York Academy of Sciences 51, 660–672 (1949).

24. T. T. Takahashi, R. W. Roberts, “In Vitro Selection of Protein and Peptide Libraries Using mRNA Display” in Methods in Molecular Biology, G. Mayer, Ed. (Humana Press, Totowa, NJ, 2009), vol. 535, chap. 17, pp. 293–314.

25. C. R. Cantor, P. R. Schimmel, Biophysical Chemistry Part III: The behavior of biologocial macromolecules (W. H. Freeman and Company, New York, 1980), 10.1016/0307-4412(81)90144-8, pp. 1371.

26. A. A. Schreier, P. R. Schimmel, Interaction of manganese with fragments, complementary fragment recombinations, and whole molecules of yeast phenylalanine specific transfer RNA. J Mol Biol 86, 601–620 (1974).

27. N. K. Pandey et al., Fluorescent protein tagging promotes phase separation and alters the aggregation pathway of huntingtin exon-1. J Biol Chem 300, 105585 (2024).

28. S. Vieweg, A. Ansaloni, Z. M. Wang, J. B. Warner, H. A. Lashuel, An Intein-based Strategy for the Production of Tag-free Huntingtin Exon 1 Proteins Enables New Insights into the Polyglutamine Dependence of Httex1 Aggregation and Fibril Formation. J Biol Chem 291, 12074–12086 (2016).

29. R. A. Azizyan et al., Establishment of Constraints on Amyloid Formation Imposed by Steric Exclusion of Globular Domains. J Mol Biol 430, 3835–3846 (2018).

30. N. Riguet et al., Nuclear and cytoplasmic huntingtin inclusions exhibit distinct biochemical composition, interactome and ultrastructural properties. Nat Commun 12, 6579 (2021).

31. J. Ko et al., Identification of distinct conformations associated with monomers and fibril assemblies of mutant huntingtin. Hum Mol Genet 27, 2330–2343 (2018).

32. D. Helmlinger et al., Progressive retinal degeneration and dysfunction in R6 Huntington’s disease mice. Hum Mol Genet 11, 3351–3359 (2002).

33. A. H. Batcha et al., Retinal dysfunction, photoreceptor protein dysregulation and neuronal remodelling in the R6/1 mouse model of Huntington’s disease. Neurobiol Dis 45, 887–896 (2012).

34. H. Xu, A. Ajayan, R. Langen, J. Chen, Pleiotropic effects of mutant huntingtin on retinopathy in two mouse models of Huntington’s disease. Neurobiol Dis 205, 106780 (2025).

35. L. D. Carter-Dawson, M. M. LaVail, Rods and cones in the mouse retina. I. Structural analysis using light and electron microscopy. J Comp Neurol 188, 245–262 (1979).

36. A. S. Falk et al., Structural Model of the Proline-Rich Domain of Huntingtin Exon-1 Fibrils. Biophysical journal 119, 2019–2028 (2020).

37. J. Ko, S. Ou, P. H. Patterson, New anti-huntingtin monoclonal antibodies: implications for huntingtin conformation and its binding proteins. Brain Research Bulletin 56, 319–329 (2001).

38. J. M. Bravo-Arredondo et al., The folding equilibrium of huntingtin exon 1 monomer depends on its polyglutamine tract. J Biol Chem 293, 19613–19623 (2018).

39. D. L. Minor, Kim, P.S., Context is a major determinant of b-sheet propensity. Nature 371, 264–267 (1994).

40. M. F. Perutz, T. Johnson, M. Suzuki, J. T. Finch, Glutamine repeats as polar zippers: their possible role in inherited neurodegenerative diseases. Proc Natl Acad Sci U S A 91, 5355–5358 (1994).

41. F. Cong, H. Varmus, Nuclear-cytoplasmic shuttling of Axin regulates subcellular localization of beta-catenin. Proc Natl Acad Sci U S A 101, 2882–2887 (2004).

42. J. S. Steffan et al., The Huntington’s disease protein interacts with p53 and CREB-binding protein and represses transcription. Proc Natl Acad Sci U S A 97, 6763–6768 (2000).

43. F. C. Nucifora, Jr., et al., Interference by huntingtin and atrophin-1 with cbp-mediated transcription leading to cellular toxicity. Science 291, 2423–2428 (2001).

44. C. H. Lo et al., Discovery of Small Molecule Inhibitors of Huntingtin Exon 1 Aggregation by FRET-Based High-Throughput Screening in Living Cells. ACS Chem Neurosci 11, 2286–2295 (2020).

45. J. G. Galaz-Montoya, S. H. Shahmoradian, K. Shen, J. Frydman, W. Chiu, Cryo-electron tomography provides topological insights into mutant huntingtin exon 1 and polyQ aggregates. Commun Biol 4, 849 (2021).

46. G. George et al., TDP43 and huntingtin Exon-1 undergo a conformationally specific interaction that strongly alters the fibril formation of both proteins. J Biol Chem 300, 107660 (2024).

47. F. J. B. Bauerlein et al., In Situ Architecture and Cellular Interactions of PolyQ Inclusions. Cell 171, 179–187 e110 (2017).

48. M. Lu et al., Live-cell super-resolution microscopy reveals a primary role for diffusion in polyglutamine-driven aggresome assembly. J Biol Chem 294, 257–268 (2019).

49. N. Kosior, B. R. Leavitt, Murine Models of Huntington’s Disease for Evaluating Therapeutics. *Methods in molecular biology (Clifton*, N.J 1780, 179–207 (2018).

50. A. Jiang, R. R. Handley, K. Lehnert, R. G. Snell, From Pathogenesis to Therapeutics: A Review of 150 Years of Huntington’s Disease Research. Int J Mol Sci 24 (2023).

51. A. Der-Sarkissian, C. C. Jao, J. Chen, R. Langen, Structural organization of alpha-synuclein fibrils studied by site-directed spin labeling. J Biol Chem 278, 37530–37535 (2003).

52. T. T. Takahashi, R. W. Roberts, In vitro selection of protein and peptide libraries using mRNA display. *Methods in molecular biology (Clifton*, N.J 535, 293–314 (2009).

53. R. W. Roberts, J. W. Szostak, RNA-peptide fusions for the *in vitro* selection of peptides and proteins. Proc Natl Acad Sci U S A 94, 12297–12302 (1997).

54. R. Liu, J. Barrick, J. W. Szostak, R. W. Roberts, Optimized Synthesis of RNA-Protein Fusions for In Vitro Protein Selection. Methods in Enzymology 317, 268–293 (2000).

55. A. Krejci, T. R. Hupp, M. Lexa, B. Vojtesek, P. Muller, Hammock: a hidden Markov model-based peptide clustering algorithm to identify protein-interaction consensus motifs in large datasets. Bioinformatics 32, 9–16 (2016).

56. T. D. Schneider, R. M. Stephens, Sequence logos: a new way to display consensus sequences. Nucleic Acids Res 18, 6097–6100 (1990).

57. G. E. Crooks, G. Hon, J. M. Chandonia, S. E. Brenner, WebLogo: a sequence logo generator. Genome Res 14, 1188–1190 (2004).

