## Supporting information for Noridomi et al 2025 for "Protofibril Binding Peptides Recognize and Inhibit Huntingtin Amyloid Formation *in vitro* and *in vivo*"

**This PDF file includes:**

Figures S1 to S7
Tables S1 to S4
SI References

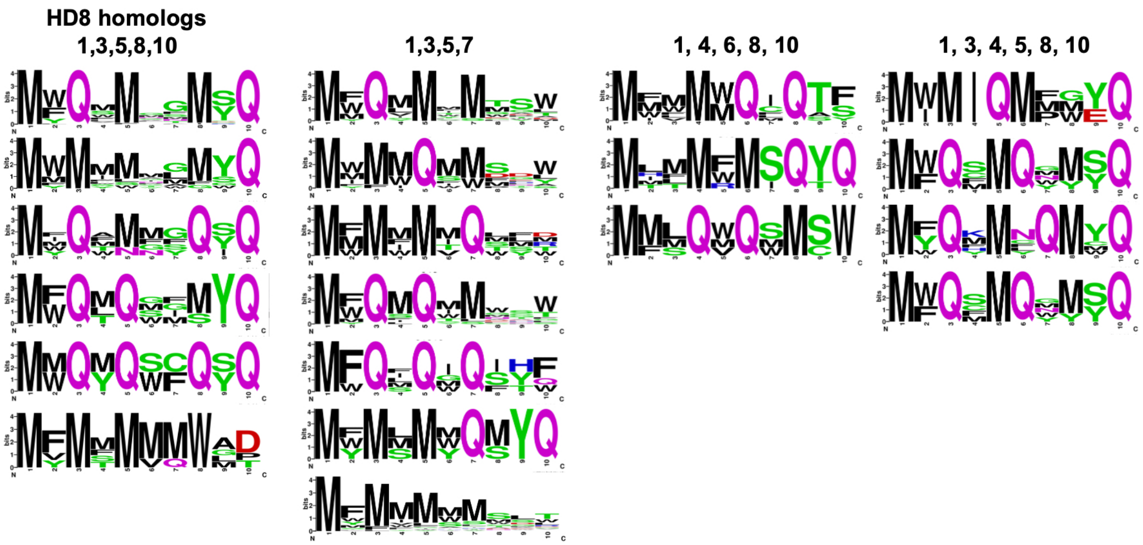

**Fig. S1.** Structure function relationships for protofibril binding peptides for Q-rich sequences. Sequence logo (1) representation of pool 8 reads containing various Q and M spacing. The position of Q or M residues is indicated above each column.

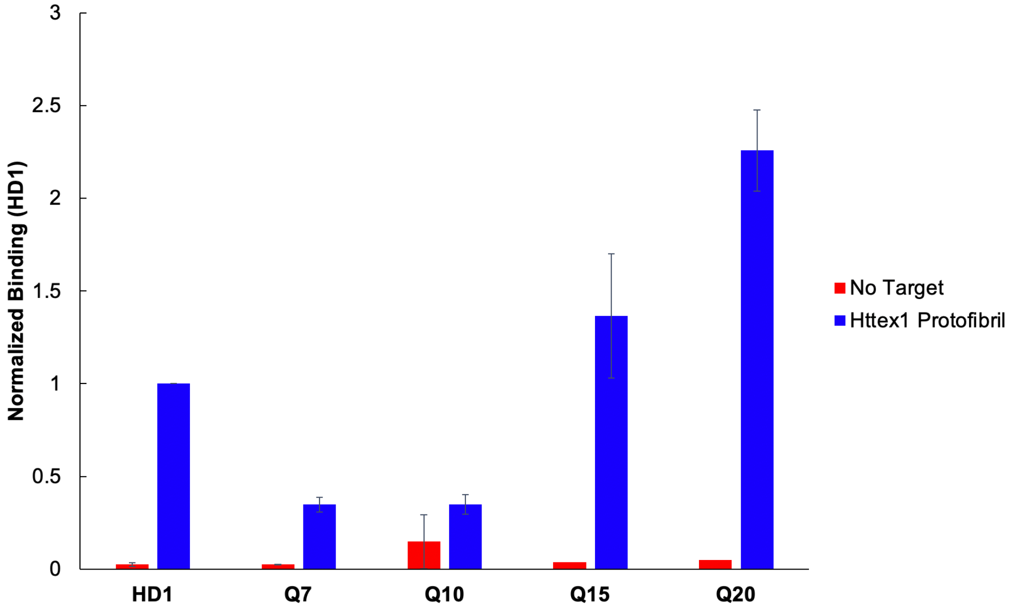

**Fig. S2**. Relative binding of HD1 and polyQ containing peptides to Httex1 protofibrils. Radioactive pulldown indicates that Q7 and Q10 peptides bind with lower affinity than HD1 and that Q15 and Q20 sequences bind with higher affinity.

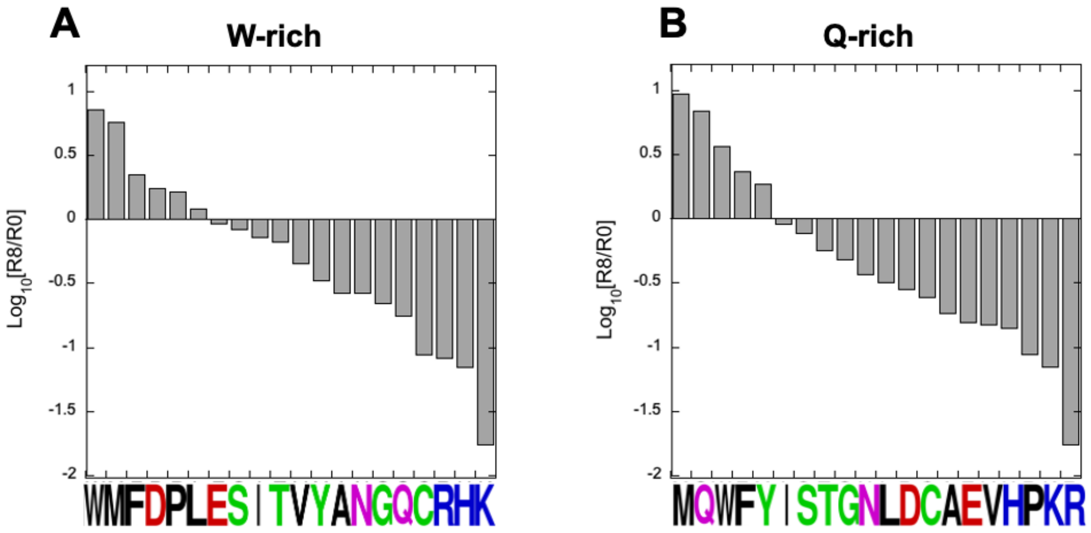

**Fig. S3.** Relative abundance of amino acids in Pool 8 vs. Pool 0 for (A) W-rich and (B) Q-rich peptides. The top ~200 W-rich and Q-rich peptide sequences in round 8 (R8) were analyzed and compared to the abundances in the naïve round 0 pool. For W-rich peptides such as HD1, hydrophobic and negatively charged residues (including proline) increase in abundance, whereas polar and positively charged residues decrease. Positively charged residues (R, H, K) show more than a 10-fold decreases compared to the naïve pool. For Q-rich sequences, methionine and glutamine have been enriched such that they comprise ~50% of the open reading frame positions, whereas proline and the positively charged residues represent only ~1.3% of the total.

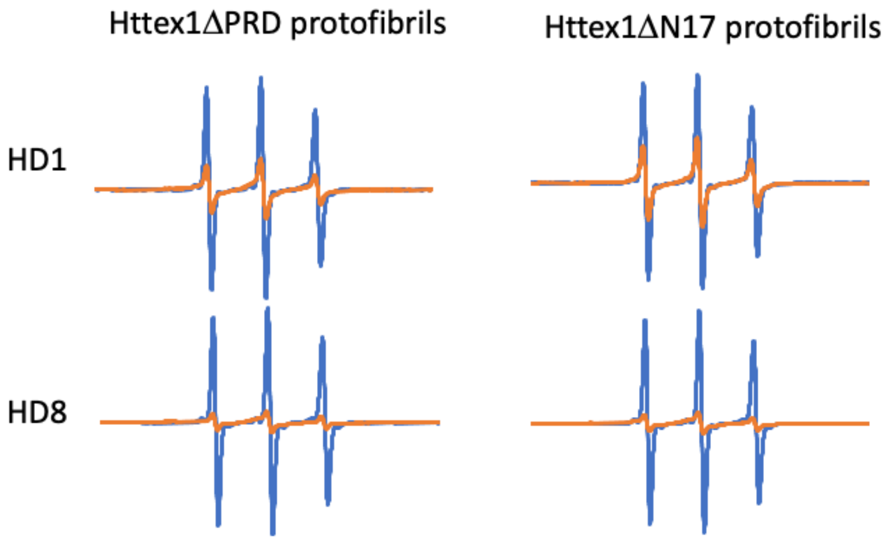

**Fig. S4**. EPR spectroscopy of spin-labeled HD1 and HD8 indicates both peptides bind specifically to the polyQ structure in Httex1 protofibrils. EPR spectra of 20 uM HD1 or HD8 are shown in the absence (blue) or presence (orange) of Httex1 protofibrils lacking the PRD (Httex1ΔPRD) or the N17 (Httex1ΔN17). The reduction in EPR amplitude indicates binding with similar amplitude loss seen in the wt Httex1 protofibrils. The data show that neither the PRD nor the N17 region are required for HD1 or HD8 binding.

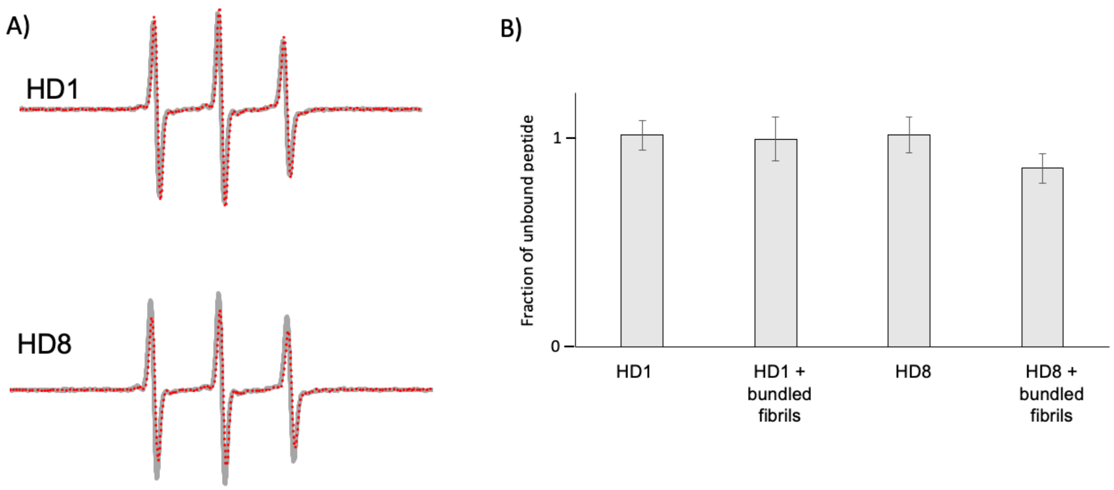

**Fig. S5**. HD1 and HD8 display little binding to bundled fibrils. A. The EPR spectra of 8 uM HD1 or HD8 in free solution (gray lines) vs. HD1 or HD8 in the presence of 8 uM bundled fibrils (red dashed lines; [Httex1 monomer] = 8 uM). B. Quantification HD1 or HD8 spectra to give the fraction of unbound peptide (+/-) bundled Httex1 fibrils. For HD1, no binding is seen to bundled fibrils, whereas for HD8 ~15% binding is observed. Six measurements were made for each condition.

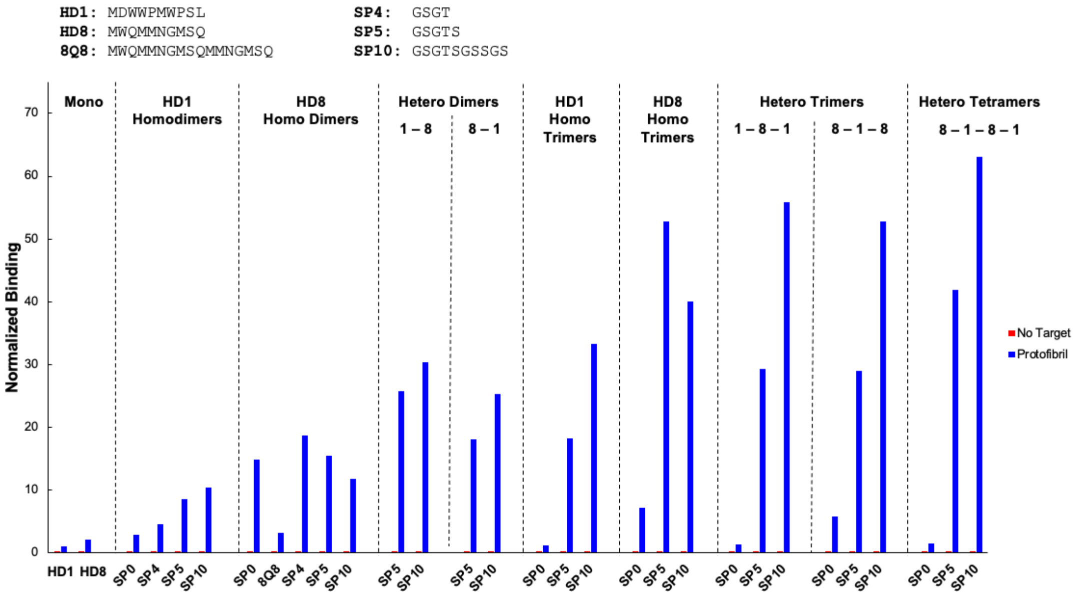

**Fig. S6**. Relative binding affinity for HD1 and HD8 monomers, dimers, trimers, and tetramers linked by various flexible spacer sequences. [$^{35}$S] labeled peptide binding was measured for pulldown using a low amount of immobilized protofibril target (35 pmol) and the binding normalized to that of HD1 under the same conditions. Generally, adding additional HD1 or HD8 peptide units attached by five-residue (SP5) or ten-residue (SP10) spacers increases binding, with the highest affinity occurring with the 10 residue flexible spacer (SP10).

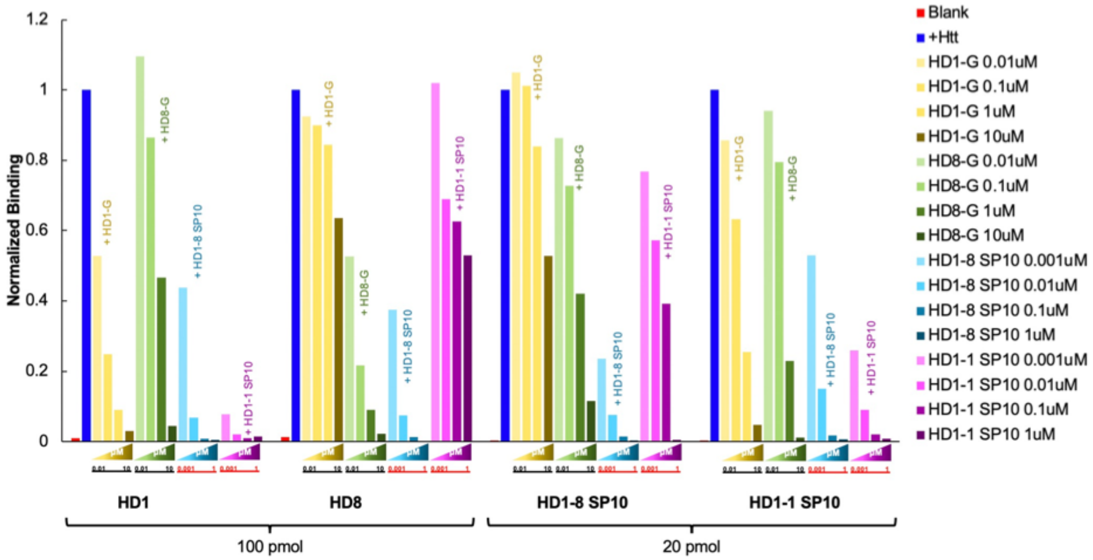

**Fig. S7**. Binding competition between labeled and unlabeled HD1, HD8, HD1-8 SP10, and HD1-1 SP10. Unlabeled peptides (HD1-G (yellow bars), HD8 (green bars), HD1-8 SP10 (blue bars), and HD1-1 SP10 (magenta bars)) were preincubated with 100 pmol or 20 pmol immobilized Httex1 protofibrils, followed by addition of [$^{35}$S] radioactively labeled HD1, HD8, HD1-8 SP10 and HD1-1 SP10. Preincubation with HD8 containing peptides (HD8 or HD1-8) reduces the binding of all the peptides tested. Preincubation with HD1 or HD1-1 reduces HD8 binding by only 40-60% indicating. HD1-8 SP10 effectively eliminates radioactive peptide binding at 1 uM concentration.

**Table S1.** Expressed Protein Sequences

| Name | Sequence |
|---|---|
| Httex1(Q46) | MATLEKLMKAFESLKSFQ$_{46}$P$_{11}$QLPQP$_{3}$QAQPLLPQPQP$_{10}$ GPAVAEEPLHRP |
| Httex1(Q46)-35R1 | MATLEKLMKAFESLKSFQ$_{17}$**R1**Q$_{28}$P$_{11}$QLPQP$_{3}$QAQPLLPQPQP$_{10}$ GPAVAEEPLHRP ∟Nitroxide |
| Httex1ΔPRD | MATLEKLMKAFESLKSFQ$_{46}$ |
| Httex1ΔN17 | MQ$_{46}$P$_{11}$QLPQP$_{3}$QAQPLLPQPQP$_{10}$GPAVAEEPLHRP |
| Httex1Q72 | MATLEKLMKAFESLKSFQ$_{72}$P$_{11}$QLPQP$_{3}$QAQPLLPQPQP$_{10}$ GPAVAEEPLHRP |

**R1 =** 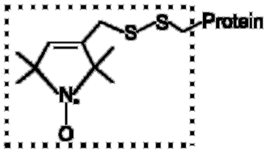

Side-chain R1 as in (2)

**Supporting Table 2.** Round 8 Sequences with more than 500 reads.

| Rank | Sequence ($MX_9$) | Reads | Name |
|---|---|---|---|
| 1 | MDWWPMWPSL | 1019561 | HD1 |
| 2 | MFFVLSWTPL | 637645 | HD5 |
| 3 | MFMMMWMSLT | 205700 | HD4 |
| 4 | MILWPMWPSL | 119133 | |
| 5 | MWQMMNGMSQ | 105044 | HD8 |
| 6 | MVFWPFWEPL | 96074 | |
| 7 | MQMWTMWEPW | 64526 | HD6 |
| 8 | MWMIQMPGYQ | 57965 | HD2 |
| 9 | MFMWDLWPDL | 56472 | |
| 10 | MDWWPMWADL | 54593 | |
| 11 | MEMWPMWPAL | 53210 | |
| 12 | MWQMMYGMSQ | 37949 | |
| 13 | MWQLMGGMYQ | 32506 | |
| 14 | MVFWELWDDW | 31920 | |
| 15 | MFMWMSMTAM | 29674 | |
| 16 | MFQSMQGMYQ | 28635 | |
| 17 | MYQWMSGMYQ | 22872 | |
| 18 | MMWMWQIQTS | 22317 | |
| 19 | MWMIMTESWM | 20908 | |
| 20 | MFMQMQMMCW | 17647 | |
| 21 | MLMWEWWPDL | 17311 | |
| 22 | MLMWAMWPDF | 16820 | |
| 23 | MMMWGMWDDW | 16056 | |
| 24 | MRWSMLLSWT | 15162 | |
| 25 | MDLWPMWESW | 14509 | HD7 |
| 26 | MIFWMMWEPL | 13783 | |
| 27 | MTMWDWWPPL | 12523 | |
| 28 | MWMMMQMQTY | 11284 | |
| 29 | MTLWPMWESW | 11142 | |
| 30 | MWQMQSMDSW | 10496 | |
| 31 | MFFWELWPPL | 10456 | |
| 32 | MFQMMGNMSQ | 10270 | |
| 33 | MFQMMMQLSW | 9836 | |
| 34 | MWMIMTMSGW | 8860 | |
| 35 | MMMMIMTMSW | 8386 | |
| 36 | MWMFMTLMYQ | 8325 | |
| 37 | MFYVLTWNPL | 8315 | |
| 38 | MAMWPMWESW | 7987 | |

| | | | |
|---|---|---|---|
| 39 | MLWWPMWESW | 7212 | |
| 40 | MIMMFMSQYQ | 7169 | |
| 41 | MEWWPMWDDW | 7165 | |
| 42 | MIMWPMWNDW | 6994 | |
| 43 | MVWMMEWADF | 6747 | |
| 44 | MRWSMSYSWA | 6737 | HD3 |
| 45 | MWQSMQYMSQ | 6150 | |
| 46 | MWPSLVVVRL | 5986 | |
| 47 | MWQMMFQMYQ | 5673 | |
| 48 | MMWWDLWPDL | 5353 | |
| 49 | MQMQWMEMSW | 5174 | |
| 50 | MMLMMQWMGQ | 5109 | |
| 51 | MWQLQGFMYQ | 4991 | |
| 52 | MDMWPMWPML | 4548 | |
| 53 | MWQMMGMMYQ | 4527 | |
| 54 | MWMYMMLGQM | 4500 | |
| 55 | MMSQWQSMSW | 4470 | |
| 56 | MFQMQWGSYQ | 4375 | |
| 57 | MFYVMSWTPL | 4340 | |
| 58 | MYMMMWQIQM | 4336 | |
| 59 | MFMSMWQMYQ | 4133 | |
| 60 | MWMMFILTPG | 4104 | |
| 61 | MWQMMYMQSW | 3939 | |
| 62 | MFQFMKGMYQ | 3860 | |
| 63 | MWMMMMQSYQ | 3833 | |
| 64 | MWMIMIMGSQ | 3812 | |
| 65 | MFMWPMWPAL | 3805 | |
| 66 | MNFWPMWDNW | 3668 | |
| 67 | MFMMMVMWQT | 3626 | |
| 68 | MYMMMMMFSQ | 3547 | |
| 69 | MMFMWQIQTS | 3513 | |
| 70 | MFQSMGLMYQ | 3453 | |
| 71 | MWQMQMMESW | 3424 | |
| 72 | MLFVLSWTPL | 3422 | |
| 73 | MYMMQFGQYQ | 3409 | |
| 74 | MVFWPLWEPL | 3314 | |
| 75 | MFFVLSWAPL | 3297 | |
| 76 | MFQWMSGMSQ | 3227 | |
| 77 | MWQSQWMNTL | 3183 | |
| 78 | MMQMMFMTSW | 3139 | |
| 79 | MTFWDFWDDL | 3112 | |

| | | |
|---|---|---|
| 80 | MFLVLSWTPL | 3106 |
| 81 | MFQFMPGMYQ | 3066 |
| 82 | MFMMMWMSSV | 2900 |
| 83 | MWQCMQNMSQ | 2705 |
| 84 | MLMMWMIQTQ | 2700 |
| 85 | MMMQMMSMCW | 2695 |
| 86 | MFQLMIMMSW | 2693 |
| 87 | MFQMMISMYQ | 2647 |
| 88 | MDWCPMWPSL | 2647 |
| 89 | MMMMWMAMYQ | 2523 |
| 90 | MLQMMGFMYQ | 2515 |
| 91 | MWLFEWWPEL | 2502 |
| 92 | MQMWPMWENW | 2486 |
| 93 | MYQMMMFMCQ | 2475 |
| 94 | MQWMMAWMSW | 2464 |
| 95 | MFYVISWTPL | 2406 |
| 96 | MMLQVQWMSW | 2295 |
| 97 | MIQMNFYQSQ | 2274 |
| 98 | MFMFQAQTFF | 2268 |
| 99 | MWQIMWMEDW | 2246 |
| 100 | MDWWPMRPSL | 2171 |
| 101 | MYQMMNQMWQ | 2153 |
| 102 | MMQWMNFQSQ | 2128 |
| 103 | MIMIQMFMYQ | 2114 |
| 104 | MVYWDMWEPW | 2074 |
| 105 | MLWWDMWPDL | 2060 |
| 106 | MMMWMWMSST | 2052 |
| 107 | MWMAMTSGWM | 2045 |
| 108 | MWMIMSGLYL | 2043 |
| 109 | MWQMQSQGSW | 2019 |
| 110 | MFQIQFQMYF | 1995 |
| 111 | MGWWPMWPSL | 1961 |
| 112 | MTWMMEWYDF | 1947 |
| 113 | MFQLMWTMYQ | 1932 |
| 114 | MFQMMIMTGW | 1922 |
| 115 | MDWWPTWPSL | 1903 |
| 116 | MFQKMLQMYQ | 1901 |
| 117 | MDWLPMWPSL | 1890 |
| 118 | MWQSMLFMSQ | 1889 |
| 119 | MNMWNMWPDL | 1881 |
| 120 | MDWRPMWPSL | 1843 |

| | | |
|---|---|---|
| 121 | MWTWMWWPDL | 1818 |
| 122 | MWVFEFWPDL | 1810 |
| 123 | MYQYMNQMYQ | 1809 |
| 124 | MFQMQSQSYW | 1789 |
| 125 | MIWWDLWEPF | 1788 |
| 126 | MIFWPLWPRL | 1768 |
| 127 | MWMMQSMSFA | 1756 |
| 128 | MYQMQIMWSG | 1755 |
| 129 | MYWMMQYQSQ | 1746 |
| 130 | MIWWPMWPEL | 1742 |
| 131 | MMFWAMWPPM | 1719 |
| 132 | MVWWPMWQEL | 1716 |
| 133 | MFMMMWMSST | 1716 |
| 134 | MWMQMGLMYQ | 1716 |
| 135 | MDWWPIWPSL | 1698 |
| 136 | MIFWEMWPSW | 1678 |
| 137 | MWQFMGSQIQ | 1666 |
| 138 | MWMMMTMNQA | 1651 |
| 139 | MWMMMQCQSF | 1629 |
| 140 | MDLWPMWPSL | 1615 |
| 141 | MFQFQTMMPW | 1607 |
| 142 | MWMMMMMSDV | 1597 |
| 143 | MWQFMATMSQ | 1591 |
| 144 | MWMIMIMFGP | 1588 |
| 145 | MFFALSWTPL | 1579 |
| 146 | MWQWMGTMSQ | 1568 |
| 147 | MWMIQWQIQA | 1564 |
| 148 | MDWWPVWPSL | 1542 |
| 149 | MYWWPMWPSL | 1541 |
| 150 | MMWMMQWMSQ | 1522 |
| 151 | MYQVMSGMYQ | 1516 |
| 152 | MMQMQMMLCW | 1516 |
| 153 | MMWMWQIQAS | 1506 |
| 154 | MDWWPMCPSL | 1497 |
| 155 | MWMLMMMNQT | 1476 |
| 156 | MFVMWQFQTF | 1449 |
| 157 | MFMMMGLMWQ | 1448 |
| 158 | MDWWPMLPSL | 1446 |
| 159 | MWQYQSFQSQ | 1440 |
| 160 | MYMMMVMWAD | 1418 |
| 161 | MNWWPMWPSL | 1403 |

| | | |
|---|---|---|
| 162 | MDFWPLWRPL | 1402 |
| 163 | MFFVLSCTPL | 1396 |
| 164 | MQMWAMWEPW | 1393 |
| 165 | MWQVMMGMYQ | 1387 |
| 166 | MDWWPMWPPL | 1374 |
| 167 | MFTFMWQIQT | 1372 |
| 168 | MWMFMMPGVM | 1354 |
| 169 | MDWWPMWPLL | 1353 |
| 170 | MDWWTMWPSL | 1337 |
| 171 | MDWWSMWPSL | 1329 |
| 172 | MWQIQIQIHF | 1328 |
| 173 | MDWWPMWPSM | 1327 |
| 174 | MDRWPMWPSL | 1322 |
| 175 | MFMWMSFDDF | 1316 |
| 176 | MWQMMFMGPT | 1308 |
| 177 | MDWWPMWTSL | 1305 |
| 178 | MMMWQMWPLW | 1300 |
| 179 | MIMWQMWDNW | 1300 |
| 180 | MLMFMWQVQT | 1290 |
| 181 | MWQMMSMLSW | 1287 |
| 182 | MILWNFWPDL | 1281 |
| 183 | MFLMRIWFDG | 1270 |
| 184 | MFFVLSLTPL | 1267 |
| 185 | MDWWLMWPSL | 1254 |
| 186 | MWQMQMIMYQ | 1247 |
| 187 | MLMWDLWESW | 1239 |
| 188 | MMYMCMSMSW | 1237 |
| 189 | MEMWPLWPPL | 1237 |
| 190 | MLMWDMWESW | 1221 |
| 191 | MWIFEWWPEW | 1216 |
| 192 | MWQSMMSMYQ | 1189 |
| 193 | MVFWEMWPLW | 1185 |
| 194 | MDLWPMWEPW | 1180 |
| 195 | MWTFEWWPTL | 1175 |
| 196 | MWMYMAGMYQ | 1170 |
| 197 | MMQSMTMMSW | 1169 |
| 198 | MDWWPMWPSL | 1164 |
| 199 | MWQMMSGMSQ | 1146 |
| 200 | MDFWPMWQSW | 1124 |
| 201 | MWMLMNLMYQ | 1124 |
| 202 | MWMMQWQSDF | 1113 |

| | | |
|---|---|---|
| 203 | MYQLMMMTSW | 1105 |
| 204 | MFQMMMSMYL | 1099 |
| 205 | MWMSMPGMYQ | 1097 |
| 206 | MFFVLPWTPL | 1097 |
| 207 | MWMMLSGMYQ | 1097 |
| 208 | MWMWMTTDQM | 1092 |
| 209 | MFFVLSRTPL | 1090 |
| 210 | MWMIMYGMYQ | 1088 |
| 211 | MWMWMSMNGL | 1084 |
| 212 | MLYVLSWTPL | 1080 |
| 213 | MWMFMSFGGL | 1073 |
| 214 | MIVMWQMQYQ | 1072 |
| 215 | MWMMQSMSYY | 1070 |
| 216 | MFQMQIQSYF | 1066 |
| 217 | MFMMMFMSLT | 1065 |
| 218 | MDWWPMWPSP | 1054 |
| 219 | MFQFQMQIHF | 1045 |
| 220 | MWQFMTNMYQ | 1044 |
| 221 | MDFWPMWDNW | 1042 |
| 222 | MDWWHMWPSL | 1040 |
| 223 | MWMIMTQTFM | 1028 |
| 224 | MLTMFMSQYQ | 1027 |
| 225 | MLMWALWPDW | 1023 |
| 226 | MFMMMWMSLA | 1012 |
| 227 | MQFWPMWEPW | 1010 |
| 228 | MFAVLSWTPL | 1008 |
| 229 | MDWWPMWHSL | 996 |
| 230 | MEMWPMWPSL | 992 |
| 231 | MWQIQIMWST | 991 |
| 232 | MFMWMMFDQM | 990 |
| 233 | MFQSMMMMSQ | 981 |
| 234 | MFQMQMMAGW | 978 |
| 235 | MFQMIQSQYQ | 977 |
| 236 | MIFWQMWDDW | 976 |
| 237 | MFMMMIQMYR | 963 |
| 238 | MWMLMQGMYQ | 961 |
| 239 | MWQIQSFAYQ | 961 |
| 240 | MWSMFIVTSR | 955 |
| 241 | MMMMQMMMDW | 954 |
| 242 | MAMWPMWESW | 953 |
| 243 | MWMFMSMMQT | 950 |

| | | |
|---|---|---|
| 244 | MYMMMFMCQI | 946 |
| 245 | MFFVLSWTPR | 934 |
| 246 | MSMMMMMGWG | 927 |
| 247 | MFQSQGQFTW | 926 |
| 248 | MWQMQWMNNW | 924 |
| 249 | MLWWDMWPDV | 918 |
| 250 | MLMFMWQIQT | 916 |
| 251 | MFMWMSMTSL | 916 |
| 252 | MFQKMYQMCQ | 914 |
| 253 | MFQFMLFGQM | 913 |
| 254 | MMWMYQVQYQ | 909 |
| 255 | MFMTMWMSLT | 908 |
| 256 | MHMMFMSQYQ | 897 |
| 257 | MDWWPMWPSR | 896 |
| 258 | MFQQQLQMYW | 890 |
| 259 | MWQSMAFMSQ | 890 |
| 260 | MMMFMMQLSW | 889 |
| 261 | MFQMMWMTTA | 879 |
| 262 | MMMMSMWMGQ | 876 |
| 263 | MFMLMISNGR | 866 |
| 264 | MVMFMMMWAD | 864 |
| 265 | MFFVLSWTLL | 863 |
| 266 | MIWWELWPMM | 859 |
| 267 | MFQMQMLSHF | 855 |
| 268 | MYQLMNGMSQ | 852 |
| 269 | MFFVLSWMPL | 851 |
| 270 | MFMMMRMSLT | 835 |
| 271 | MWEMWPSLTF | 830 |
| 272 | MSFVLSWTPL | 830 |
| 273 | MMQYMMMWAD | 826 |
| 274 | MLMWAMWPDL | 820 |
| 275 | MVWWDLWDDL | 819 |
| 276 | MFQMMQMYYQ | 818 |
| 277 | MTWWPLWESM | 818 |
| 278 | MFSVLSWTPL | 816 |
| 279 | MFQMQMMGCW | 816 |
| 280 | MFQMQTMWAT | 816 |
| 281 | MFLWAWWPEL | 813 |
| 282 | MYMYMMMNEL | 810 |
| 283 | MWQIMANMSQ | 808 |
| 284 | MMWQMMWMDA | 807 |

| | | |
|---|---|---|
| 285 | MWQMMYMSGL | 805 |
| 286 | MWMSMTMYDW | 803 |
| 287 | MFMMMMMSWG | 798 |
| 288 | MWMMQIMFES | 794 |
| 289 | MWFMMWQSQT | 793 |
| 290 | MFFVLSWTPP | 792 |
| 291 | MNFWPMWEWT | 791 |
| 292 | MIMMMSQWQT | 778 |
| 293 | MMMWMSLTGS | 772 |
| 294 | MDLWEWWPDL | 767 |
| 295 | MFFVLSWTPM | 765 |
| 296 | MWMIMYMTEQ | 765 |
| 297 | MFLIFSLDGR | 762 |
| 298 | MWMMMQMSHF | 755 |
| 299 | MFQTQSMMYQ | 754 |
| 300 | MMMMWMSMHF | 754 |
| 301 | MMLQWQSMSW | 753 |
| 302 | MPMWPMWECW | 746 |
| 303 | MFMMMWISLT | 745 |
| 304 | MWMMMWQDQT | 744 |
| 305 | MIMIMMMMGW | 744 |
| 306 | MDCWPMWPSL | 743 |
| 307 | MFMIMWMSLT | 741 |
| 308 | MFQMQWMCQS | 741 |
| 309 | MWMMQFQSYQ | 741 |
| 310 | MISWGFLIVL | 739 |
| 311 | MFFVLSWTPL | 738 |
| 312 | MMLMTMWMSW | 731 |
| 313 | MWNMFYITAG | 730 |
| 314 | MFFVLSWTQL | 728 |
| 315 | MLMMMWMSLT | 726 |
| 316 | MFQAMMGQYQ | 722 |
| 317 | MYMIMMMMSW | 718 |
| 318 | MFQMMLMWST | 717 |
| 319 | MWQCMLGMSQ | 717 |
| 320 | MWQQMSMSWM | 714 |
| 321 | MWMFMMMGEM | 713 |
| 322 | MFFVLLWTPL | 704 |
| 323 | MWMMMMSWST | 704 |
| 324 | MWMIQMMWEQ | 699 |
| 325 | MMFMRMSQYQ | 696 |

| | | |
|---|---|---|
| 326 | MWMYMSFDGF | 693 |
| 327 | MFMSMQMWTT | 686 |
| 328 | MYMFMMFQTF | 684 |
| 329 | MYMLMYQMYQ | 684 |
| 330 | MFFVSSWTPL | 683 |
| 331 | MHFWPMWPQL | 682 |
| 332 | MTLMWMSQYQ | 677 |
| 333 | MSMMMSMMSW | 677 |
| 334 | MMMMQMMMGW | 673 |
| 335 | MYVQWQMQYQ | 673 |
| 336 | MFMLMWQVQA | 667 |
| 337 | MYQHMNQMYQ | 666 |
| 338 | MIFWTLWPPL | 666 |
| 339 | MLWWPQWESL | 665 |
| 340 | MFMWMTSSQM | 663 |
| 341 | MFQSQWMGLT | 662 |
| 342 | MWQMMHGMSQ | 662 |
| 343 | MDWWPMWSSL | 658 |
| 344 | MYMMMWMSLT | 658 |
| 345 | MFMMMWMSLT | 653 |
| 346 | MWQFMQWMSQ | 653 |
| 347 | MIFWELWPAF | 645 |
| 348 | MFMMMWMGLT | 640 |
| 349 | MWMMQFMSDW | 640 |
| 350 | MMFMMMFQTS | 636 |
| 351 | MWQLQIMMDW | 635 |
| 352 | MFQWMMAMYQ | 634 |
| 353 | MIWWDLWPEF | 633 |
| 354 | MWMMMTSAYL | 632 |
| 355 | MFMLMWMIQT | 631 |
| 356 | MFFVLSWTSL | 630 |
| 357 | MRIWMKLSLT | 628 |
| 358 | MIFWEMWESW | 625 |
| 359 | MFFWPMWPTL | 619 |
| 360 | MFFILSWTPL | 619 |
| 361 | MNWWPSWPTL | 619 |
| 362 | MLFWDMWPDL | 618 |
| 363 | MWQLMVMMSW | 616 |
| 364 | MWIVILIFRT | 615 |
| 365 | MDWWPMWLSL | 614 |
| 366 | MVWWPMWPSL | 612 |

| | | |
|---|---|---|
| 367 | MLMWDLWPDL | 609 |
| 368 | MEMWPLWPML | 607 |
| 369 | MLMWPMWDDW | 606 |
| 370 | MMQMQWCQYQ | 598 |
| 371 | MFQMQYMWGS | 598 |
| 372 | MWMMMAMSLT | 596 |
| 373 | MFMWMMMLDV | 596 |
| 374 | MWQYMNGMGQ | 591 |
| 375 | MFIMMWMSLT | 586 |
| 376 | MILWPMWSSL | 582 |
| 377 | MFMMQWMSTL | 582 |
| 378 | MMFMTMSMWA | 579 |
| 379 | MWQMMFGMRQ | 578 |
| 380 | MFMSMMMWGT | 569 |
| 381 | MDSWEWWPDL | 569 |
| 382 | MDWWPLWPSL | 565 |
| 383 | MFMTMMMWMP | 561 |
| 384 | MWMKMSTMYQ | 557 |
| 385 | MFMMMQLQTF | 552 |
| 386 | MFQSMLSMSQ | 551 |
| 387 | MWSQMQWQTL | 550 |
| 388 | MFQFMCQMMQ | 550 |
| 389 | MFLMMQIQTF | 547 |
| 390 | MFMMFMWMDA | 545 |
| 391 | MMWMSQSMTY | 545 |
| 392 | MFQLQWQSYQ | 545 |
| 393 | MFFVLSWTTL | 540 |
| 394 | MLWWPMWADV | 537 |
| 395 | MDWWPMWETW | 536 |
| 396 | MFMIMWMVQT | 532 |
| 397 | MFDWVWWPSL | 532 |
| 398 | MYMMMAMMDW | 531 |
| 399 | MFQSFGSMYQ | 528 |
| 400 | MWMMMVMLQT | 528 |
| 401 | MVFVLSWTPL | 527 |
| 402 | MFFVFSWTPL | 527 |
| 403 | MWQMMNGISQ | 527 |
| 404 | MIMMWQCQTF | 526 |
| 405 | MYMIQMMSHY | 525 |
| 406 | MMMWMVQSMT | 524 |
| 407 | MWQVMTMSWM | 523 |

| | | |
|---|---|---|
| 408 | MFMIYSYAGR | 523 |
| 409 | MFQVMYMTSW | 520 |
| 410 | MFMFMIMWEL | 519 |
| 411 | MFMMMMQWLD | 518 |
| 412 | MFMMMSMFSW | 517 |
| 413 | MYMIMTMYSW | 517 |
| 414 | MWMMQMMFGL | 514 |
| 415 | MMWMFMSSGV | 512 |
| 416 | MWMNMWMSDW | 511 |
| 417 | MMQMQMSWTW | 510 |
| 418 | MYQTMMGQYQ | 507 |
| 419 | MYQWMMMMDQ | 506 |

**Table S3.** Amino acid sequences of peptides evaluated in radioligand binding assays**.**

| Name | Sequence |
|---|---|
| HD1 | MDWWPMWPSLGSGTSGSS |
| HD2 | MWMIQMPGYQGSGTSGSS |
| HD3 | MRWSMSYSWAGSGTSGSS |
| HD4 | MFMMMWMSLTGSGTSGSS |
| HD5 | MFFVLSWTPLGSGTSGSS |
| HD6 | MQMWTMWEPWGSGTSGSS |
| HD7 | MDLWPMWESWGSGTSGSS |
| HD8 | MWQMMNGMSQGSGTSGSS |
| HD1 MM | MMDWWPMWPSLGSGTSGSS |
| HD1 M1A | MADWWPMWPSLGSGTSGSS |
| HD1 D2A | MAWWPMWPSLGSGTSGSS |
| HD1 W3A | MDAWPMWPSLGSGTSGSS |
| HD1 W4A | MDWAPMWPSLGSGTSGSS |
| HD1 P5A | MDWWAMWPSLGSGTSGSS |
| HD1 M6A | MDWWPAWPSLGSGTSGSS |
| HD1 W7A | MDWWPMAPSLGSGTSGSS |
| HD1 P8A | MDWWPMWASLGSGTSGSS |
| HD1 S9A | MDWWPMWPALGSGTSGSS |
| HD1 L10A | MDWWPMWPSAGSGTSGSS |
| HD1 SC | MSWMLPWPWDGSGTSGSS |
| HD8 MM | MMWQMMNGMSQGSGTSGSS |
| HD8 M1A | MAWQMMNGMSQGSGTSGSS |
| HD8 W2A | MAQMMNGMSQGSGTSGSS |
| HD8 Q3A | MWAMMNGMSQGSGTSGSS |
| HD8 M4A | MWQAMNGMSQGSGTSGSS |
| HD8 M5A | MWQMANGMSQGSGTSGSS |
| HD8 N6A | MWQMMAGMSQGSGTSGSS |
| HD8 G7A | MWQMMNAMSQGSGTSGSS |
| HD8 M8A | MWQMMNGASQGSGTSGSS |
| HD8 S9A | MWQMMNGMAQGSGTSGSS |
| HD8 Q10A | MWQMMNGMSAGSGTSGSS |
| HD8 SC | MQNMSWGMQMGSGTSGSS |
| QBP1 | MSNWKWWPGIFDGSGTSGSS |
| Q7 | MQQQQQQQGSGTSGGS |
| Q10 | MQQQQQQQQQQGSGTSGGS |
| Q15 | MQQQQQQQQQQQQQQQGSGTSGGS |
| Q20 | MQQQQQQQQQQQQQQQQQQQQGSGTSGGS |
| HD1-1 SP0 | MDWWPMWPSLMDWWPMWPSLGSGTSGSS |
| HD1-1 SP4 | MDWWPMWPSLGSGTMDWWPMWPSLGSGTSGSS |
| HD1-1 SP5 | MDWWPMWPSLGSGTSMDWWPMWPSLGSGTSGSS |
| HD1-1 SP10 | MDWWPMWPSLGSGTSGSSGSMDWWPMWPSLGSGTSGSS |
| HD8-8 SP0 | MWQMMNGMSQMWQMMNGMSQGSGTSGSS |
| 8Q8 | MWQMMNGMSQMMNGMSQGSGTSGSS |
| HD8-8 SP0 | MWQMMNGMSQMWQMMNGMSQGSGTSGSS |

| | |
|---|---|
| HD8-8 SP4 | MWQMMNGMSQGSGTMWQMMNGMSQGSGTSGSS |
| HD8-8 SP5 | MWQMMNGMSQGSGTSMWQMMNGMSQGSGTSGSS |
| HD8-8 SP10 | MWQMMNGMSQGSGTSGSSGSMWQMMNGMSQGSGTSGSS |
| HD1-8 SP5 | MDWWPMWPSLGSGTSMWQMMNGMSQGSGTSGSS |
| HD1-8 SP10 | MDWWPMWPSLGSGTSGSSGSMWQMMNGMSQGSGTSGSS |
| HD8-1 SP5 | MWQMMNGMSQGSGTSMDWWPMWPSLGSGTSGSS |
| HD8-1 SP10 | MWQMMNGMSQGSGTSGSSGSMDWWPMWPSLGSGTSGSS |
| HD1-1-1 SP0 | MDWWPMWPSLMDWWPMWPSLMDWWPMWPSLGSGTSGSS |
| HD1-1-1 SP5 | MDWWPMWPSLGSGTSMDWWPMWPSLGSGTSMDWWPMWPSL<br>GSGTSGSS |
| HD1-1-1 SP10 | MDWWPMWPSLGSGTSGSSGSMDWWPMWPSLGSGTSGSSGS<br>MDWWPMWPSLGSGTSGSS |
| HD8-8-8 SP0 | MWQMMNGMSQMWQMMNGMSQMWQMMNGMSQGSGTSGSS |
| HD8-8-8 SP5 | MWQMMNGMSQGSGTSMWQMMNGMSQGSGTSMWQMMNGMSQ<br>GSGTSGSS |
| HD8-8-8 SP10 | MWQMMNGMSQGSGTSGSSGSMWQMMNGMSQGSGTSGSSGS<br>MWQMMNGMSQGSGTSGSS |
| HD1-8-1 SP0 | MDWWPMWPSLMWQMMNGMSQMDWWPMWPSLGSGTSGSS |
| HD1-8-1 SP5 | MDWWPMWPSLGSGTSMWQMMNGMSQGSGTSMDWWPMWPSL<br>GSGTSGSS |
| HD1-8-1 SP10 | MDWWPMWPSLGSGTSGSSGSMWQMMNGMSQGSGTSGSSGS<br>MDWWPMWPSLGSGTSGSS |
| HD8-1-8 SP0 | MWQMMNGMSQMDWWPMWPSLMWQMMNGMSQGSGTSGSS |
| HD8-1-8 SP5 | MWQMMNGMSQGSGTSMDWWPMWPSLGSGTSMWQMMNGMSQ<br>GSGTSGSS |
| HD8-1-8-1 SP0 | MWQMMNGMSQMDWWPMWPSLMWQMMNGMSQMDWWPMWPSL |
| HD8-1-8-1 SP5 | MWQMMNGMSQGSGTSMDWWPMWPSLGSGTSMWQMMNGMSQ<br>GSGTSMDWWPMWPSLGSGTSGSS |
| HD8-1-8-1 SP10 | MWQMMNGMSQGSGTSGSSGSMDWWPMWPSLGSGTSGSSGS<br>MWQMMNGMSQGSGTSGSSGSMDWWPMWPSLGSGTSGSS |

SP4 = GSGT
SP5 = GSGTS
SP10 = GSGTSGSSGS
Linker = GSGTSGSS
HD1 = MDWWPMWPSL
HD8 = MWQMMNGMSQ
QBP1 MSNWKWWPGIFD

**Table S4.** Amino acid sequence and structure of synthetic peptides.

| **Name** | **Sequence** |
|---|---|
| HD1 | MDWWPMWPSL-$NH_2$ |
| HD8 | MWQMMNGMSQ-$NH_2$ |
| HD1-G | MDWWPMWPSLG-$NH_2$ |
| HD8-G | MWQMMNGMSQG-$NH_2$ |
| HD8-1 SP10 | MWQMMNGMSQGSGTSGSSGSMDWWPMWPSL-$NH_2$ |
| HD1-8 SP10 | MDWWPMWPSLGSGTSGSSGSMWQMMNGMSQG-$NH_2$ |
| HD8-1 SP10 Fl | MWQMMNGMSQGSGTSGSSGSMDWWPMWPSLGGC-$NH_2$ <br> └Fl |

Fl = Fluorescein

### SI References

1. G. E. Crooks, G. Hon, J. M. Chandonia, S. E. Brenner, WebLogo: a sequence logo generator. *Genome Res* **14**, 1188-1190 (2004).
2. A. Der-Sarkissian, C. C. Jao, J. Chen, R. Langen, Structural organization of alpha-synuclein fibrils studied by site-directed spin labeling. *J Biol Chem* **278**, 37530-37535 (2003).
